# osl-dynamics: A toolbox for modelling fast dynamic brain activity

**DOI:** 10.1101/2023.08.07.549346

**Authors:** C. Gohil, R. Huang, E. Roberts, M.W.J. van Es, A.J. Quinn, D. Vidaurre, M.W. Woolrich

## Abstract

Neural activity contains rich spatio-temporal structure that corresponds to cognition. This includes oscillatory bursting and dynamic activity that span across networks of brain regions, all of which can occur on timescales of a tens of milliseconds. While these processes can be accessed through brain recordings and imaging, modelling them presents methodological challenges due to their fast and transient nature. Furthermore, the exact timing and duration of interesting cognitive events is often a priori unknown. Here we present the OHBA Software Library Dynamics Toolbox (`osl-dynamics`), a Python-based package that can identify and describe recurrent dynamics in functional neuroimaging data on timescales as fast as tens of milliseconds. At its core are machine learning generative models that are able to adapt to the data and learn the timing, as well as the spatial and spectral characteristics, of brain activity with few assumptions. `osl-dynamics` incorporates state-of-the-art approaches that can be, and have been, used to elucidate brain dynamics in a wide range of data types, including magneto/electroencephalography, functional magnetic resonance imaging, invasive local field potential recordings and electrocorticography. It also provides novel summary measures of brain dynamics that can be used to inform our understanding of cognition, behaviour and disease. We hope `osl-dynamics` will further our understanding of brain function, through its ability to enhance the modelling of fast dynamic processes.

**Highlights:**

- An open-source toolbox for identifying and describing brain dynamics in neuroimaging data on fast timescales.
- Includes visualisation and quantification of oscillatory bursting and network dynamics.
- Provides novel summary measures and group analysis tools of brain dynamics that can be used to inform our understanding of cognition, behaviour and disease.
- Implemented in Python and makes use of `TensorFlow`.
- Includes comprehensive documentation and tutorials.

## 1 Introduction

There is growing evidence for the importance of oscillatory activity in brain function [1]. Neural oscillations have been linked to cognitive processes, such as information encoding and processing, as well as attention [2] and distinct oscillatory activity has been observed in different states of consciousness [3]. Furthermore, the synchronisation of neural oscillations has been proposed as a mechanism for communication [4]. Neural oscillations have also been a useful tool for understanding brain dysfunction; for example, changes have been observed in the oscillatory activity of diseased and healthy cohorts [5].

An aspect of neural oscillations that remains to be fully understood is its dynamic nature, particularly at fast timescales [6]. Recently, it has been proposed that neuronal populations exhibit short *bursts* of oscillatory activity on timescales of 100 ms [7, 8, 9] rather than the classical view of ongoing oscillations that are modulated in amplitude. This has important implications for how we should be modelling oscillatory activity changes in cognition and disease [10, 11, 12]. Unfortunately, the methods available to detect bursts in oscillatory data are limited, often requiring arbitrary choices for parameters relating to the frequency content, amplitude and duration [13]. These choices can significantly impact the conclusions reached by such analyses.

Furthermore, oscillatory bursting is not isolated to individual brain regions. It has been shown that bursting can occur across cortical networks [9], and there are bursts of coherent activity in networks that lasts on the order of 50-100 ms in both resting-state [14] and in task [15]. Precise knowledge of these fast network dynamics is a valuable insight that can help us understand cognitive processes; for example, the dynamics of specific functional resting-state networks have been linked to memory replay (a `≤` 50 ms process that occurs in memory consolidation) [16]. Changes in the dynamics of functional networks have also been shown to be predictive of behavioural traits [17] and disease [18, 19, 20, 21, 22, 23]. The key barrier that prevents us from fully utilising a network perspective is that the accurate estimation of dynamic functional networks is challenging. This is in part due to the timing and duration of interesting cognitive events, and the corresponding activity in functional networks, not being known. Consequently, we need to rely on methods that can adapt to the data and automatically identify when networks activate.

Here, we present the OHBA Software Library Dynamics Toolbox (`osl-dynamics`), a Python package that meets two far-reaching methodological challenges that limit the field of cognitive neuroscience: burst detection and the identification of dynamic functional brain networks. It does so by deploying data driven generative models that have a proven ability to adapt to the data from a wide range of imaging modalities, and can learn the spatio-temporal characteristics of brain activity, with few assumptions and at fast timescales [24, 25, 26].

In applications for burst detection, `osl-dynamics` can automatically detect oscillatory bursts without the need to specify the frequencies, amplitude threshold or duration of the bursts. This allows `osl-dynamics` to answer questions such as: when do oscillatory bursts occur; what is their oscillatory frequency; and what are their characteristic features (e.g. average lifetime, interval, occurrence and amplitude)?

In the detection of dynamic functional brain networks, `osl-dynamics` can automatically detect network dynamics at fast timescales with few assumptions. This allows `osl-dynamics` to answer questions such as: what large-scale functional networks do individuals or groups exhibit; when do these functional networks activate and what are their characteristic dynamics; what functional networks activate in response to a task; do individuals differ in their functional network activations? On top of this, `osl-dynamics` can characterise functional networks from a more conventional, static (time-averaged), perspective using the same methodology where appropriate as the dynamic methods.

Here, we will illustrate the use of `osl-dynamics` using publicly available magnetoencephalography (MEG) datasets. However, we emphasise that the scope of the toolbox extends well beyond MEG, containing approaches that can be used, and have been used, to elucidate network and oscillatory dynamics in a range of applications that include electroencephalography [27, 28], functional magnetic resonance imaging (fMRI) [25, 30], invasive local field potential recordings [11, 29] and electrocorticography [31].

## 2 Methods

### 2.1 Generative Models

In the study of dynamics, we are often interested in the properties of a time series, such as power spectral density (PSD), mean, covariance, etc., at a given time point. A common heuristic approach for calculating this is to use a sliding window. However, this approach only utilises a short window around the time point of interest and suffers from a tradeoff between the temporal precision of dynamics and an accurate estimation of the properties (via a sufficiently large window). In [32], it was shown that this approach is inadequate for studying fast changes in functional connectivity. In `osl-dynamics`, we adopt an alternative approach based on *generative models* [33]. These are models that learn the probability distribution of the training data. In this report, we will focus on two generative models for time series data: the Hidden Markov Model (HMM) [34] and Dynamic Network Modes (DyNeMo) [26]. Both of these models (discussed further below) incorporate an underlying dynamic latent variable in the generative process. The objective during training is to learn the most likely latent variables to have generated the observed time series (we minimise the *variational free energy* ^1^ [35]). In doing this, the model can identify non-contiguous segments of the time series that share the same latent variable. Pooling this information leads to more robust estimates of the local properties of the data.

The generative model for the HMM (shown in Figure 1A) is

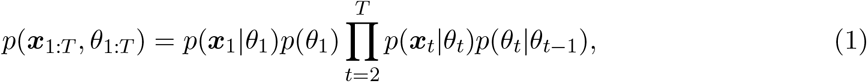

where ***θ***_*t*_ ∈ *{*1, …, *K}* is the latent state at time *t, K* is the number of states and *x*_*t*_ is the generated data. *p*(*x*_*t*_|*θ*_*t*_) is the *observation model*. Here, we use

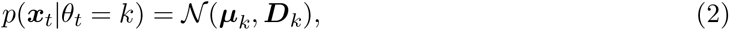

where ***µ***_*k*_ ∈ {***µ***_1_, …, ***µ***_*K*_ *}* is a state mean and ***D***_*k*_ ∈ {***D***_1_, …, D_*K*_*}* is a state covariance. Dynamics in the time series are generated through state switching, which is characterised by the transition probability *p*(*θ*_*t*_|*θ*_*t-*1_). Each pairwise state transition forms the *transition probability matrix*, [34]

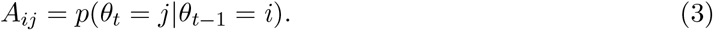

`osl-dynamics` uses variational Bayesian inference [35] to learn the most likely state to have generated the observed data. This has the advantage of being able to account for uncertainty in the latent state. For more information regarding the implementation of the HMM in `osl-dynamics` see the documentation: https://osl-dynamics.readthedocs.io/en/latest/models/hmm.html. The HMM has been successfully used to study dynamics in neuroimaging data in a variety of settings [9, 11, 14, 15, 16, 18, 20, 24, 25, 30].

**Figure 1:**
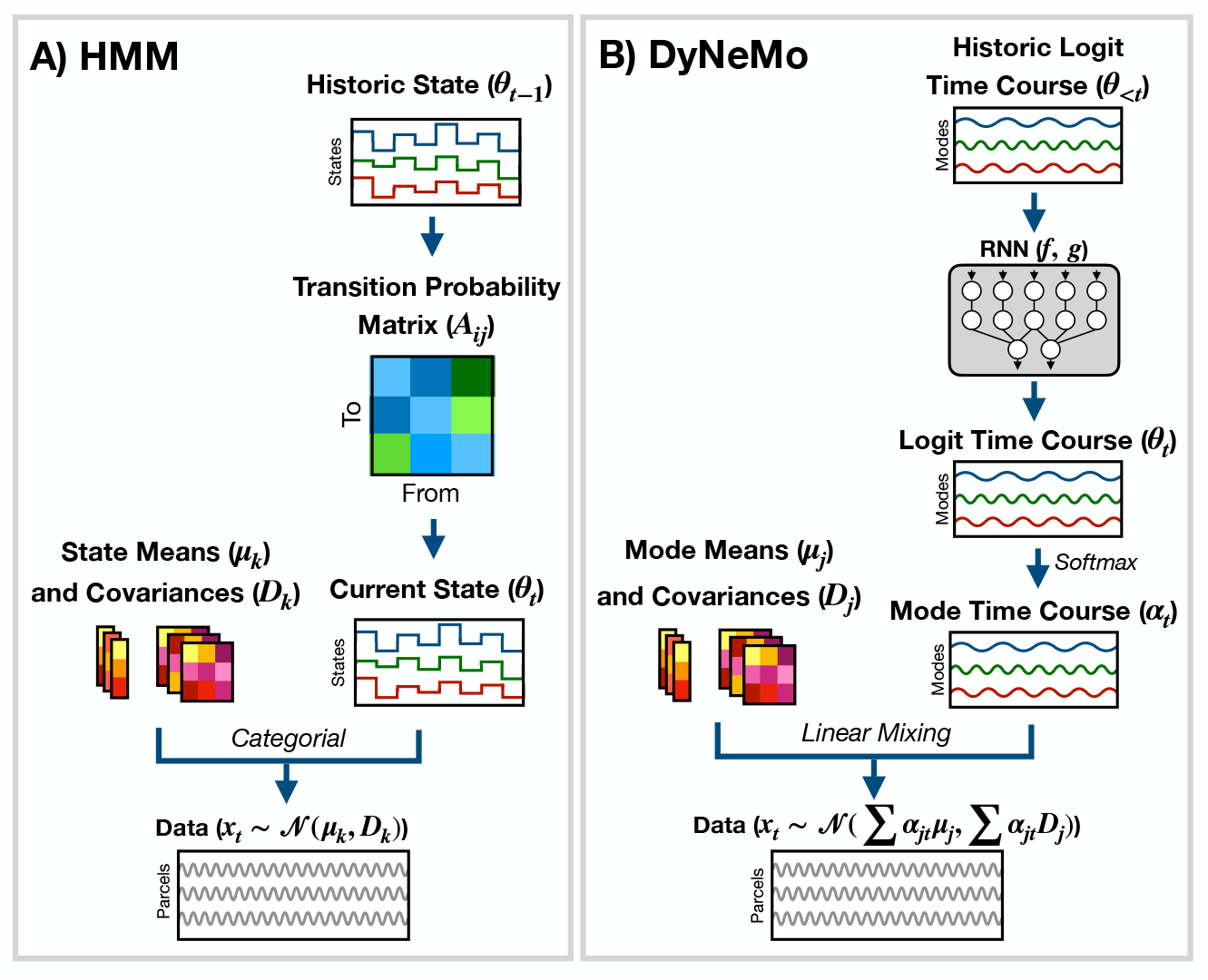
Generative models implemented in osl-dynamics. A) Hidden Markov Model (HMM) [24, 25]. Here, data is generated using a hidden state (*θ*_*t*_) and observation model, which in our case is a multivariate normal distribution parameterised by a state mean (***µ***_*k*_) and covariance (***D***_*k*_). Only one state can be active at a given time point. Dynamics are modelled via state switching using a transition probability matrix (A_*ij*_), which forecasts the probability of the current state based on the previous state. B) Dynamic Network Modes (DyNeMo) [26]. Here, the data is generated using a linear combination of *modes* (***µ***_*j*_ and D_*j*_) and dynamics are modelled using a recurrent neural network (RNN: f and g), which forecasts the probability of a particular mixing ratio (***α***_*t*_) based on a long history of previous values via the underlying logits (*θ*_*t*_).

DyNeMo is a recently proposed model that overcomes two key limitations of the HMM: the mutually exclusive states and limited memory [26]. The generative model for DyNeMo (shown in Figure 1B) is

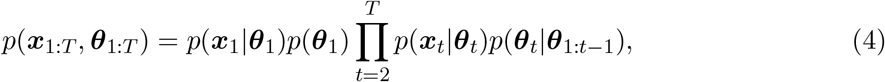

where *θ*_*t*_ is a latent vector at time t (referred to as a *logit*) and ***x***_*t*_ is the generated data. The observation model we use is

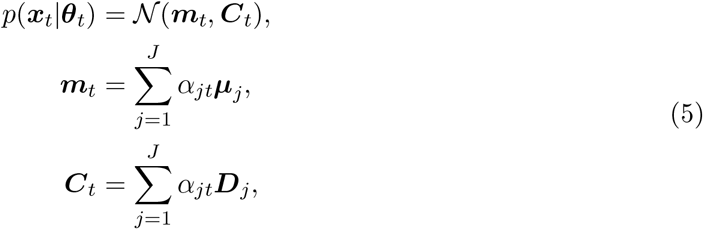

where ***µ***_*j*_ ∈ {***µ***_1_, …, ***µ***_*J*_ } is a mode mean, ***D***_*j*_ ∈ {***D***_1_, …, ***D***_*J*_ *}* is a mode covariance, *J* is the number of modes and

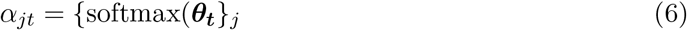

is the mixing coefficient for mode j. Dynamics in the latent vector are generated through p(***θ***_*t*_|***θ***_1:*t-*1_), which is a distribution parameterised using a recurrent neural network [36]. Specifically,

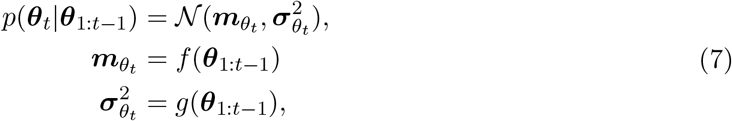

where f and g are calculated using a recurrent neural network. `osl-dynamics` uses *amortised* variational Bayesian inference [37] to learn the most likely latent vector to have generated the observed data. This is a highly efficient inference scheme that is scalable to large datasets. For more information regarding the implementation of DyNeMo in `osl-dynamics` see the documentation: https://osl-dynamics.readthedocs.io/en/latest/models/dynemo.html.

Once trained, both models reveal a dynamic latent description of the training data. For the HMM, the latent description is a hidden state time course^2^, which is the most likely state inferred at each time point in the training data. For DyNeMo, it is a mode time course, which is the mixing coefficient time series for each mode inferred from the training data. We will discuss in Sections 2.5.1 and 2.5.2 how these latent descriptions can be used to summarise dynamics in the training data.

### 2.2 Datasets

We make use of two publicly available datasets:

- **CTF rest MEG dataset**. This contains resting-state (eyes open) MEG data collected using a 275-channel CTF scanner. This dataset contains 5 minute recordings from 65 healthy participants. It was collected at Nottingham University, UK as part of the MEGUK partnership [38].
- **Elekta task MEG dataset**. This contains MEG data recorded during a visual perception task [39]. 6 runs from 19 healthy participants were recorded using an Elekta Neuromag Vectorview 306 scanner. This dataset was collected at Cambridge University, UK.

### 2.3 Preprocessing and Source Reconstruction

The steps involved in estimating source data from an MEG recording are shown in Figure 2. This part of the pipeline can be performed with the OHBA Software Library (OSL) [40, 41], which is a separate Python package for M/EEG analysis. The exact steps applied to the raw data for each dataset were:

1. MaxFilter (only applied to the Elekta dataset).
2. Bandpass filter 0.5-125 Hz.
3. Notch filter 50 Hz and 100 Hz.
4. Downsample to 250 Hz.
5. Automated bad segment removal and bad channel detection.^3^
6. Automated ICA cleaning using the correlation the EOG/ECG channel to select artefact components.^4^
7. Coregistration (using polhemus headshape points/fiducials and a structural MRI).
8. Bandpass filter 1-45 Hz.
9. Linearly Constrained Minimum Variance (LCMV) beamformer.
10. Parcellate to regions of interest. In this work, we used 38 parcels^5^.
11. Symmetric orthogonalisation (to correct source leakage [42]).
12. Dipole sign flipping (to align the sign of parcel time courses across subjects/runs).^6^
13. Downsample to 100 Hz (only included in the burst detection pipeline).

**Figure 2:**
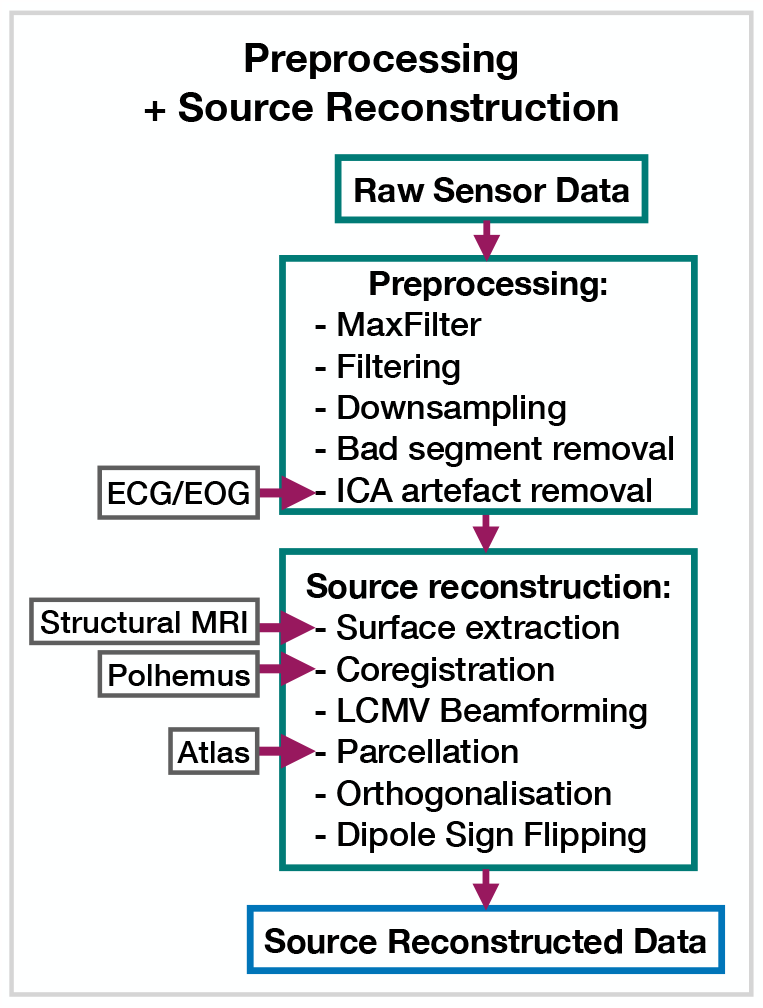
Preprocessing and source reconstruction. First, the sensor-level recordings are cleaned using standard signal processing techniques. This includes filtering, downsampling and artefact removal. Following this, the sensor-level recordings are used to estimate source activity using a beamformer. Finally, we parcellate the data and perform corrections (orthogonalisation and dipole sign flipping). Acronyms: electrocardiogram (ECG), electrooculogram (EOG), independent component analysis (ICA), Linearly Constrained Minimum Variance (LCMV). These steps can be performed with the OHBA Software Library: https://github.com/OHBA-analysis/osl.

These preprocessing steps have been found to work well for a wide variety of datasets when studying dynamics. The scripts used for preprocessing and source reconstruction can be found here: https://github.com/OHBA-analysis/osl-dynamics/tree/main/examples/toolbox_paper.

### 2.4 Data Preparation

We usually prepare the source data before training a model. The data preparation can be different depending on what aspect of the data we are interested in studying.

**Amplitude Envelope (AE)** If we are interested in studying dynamics in the amplitude of oscillations, we can train a model on AE data. Here, we typically bandpass filter a frequency range of interest and calculate an AE using the absolute value of a Hilbert transform. Figure 3B shows what happens when we calculate the AE of oscillatory data. We can see the AE data tracks changes in the amplitude of oscillations.

**Figure 3:**
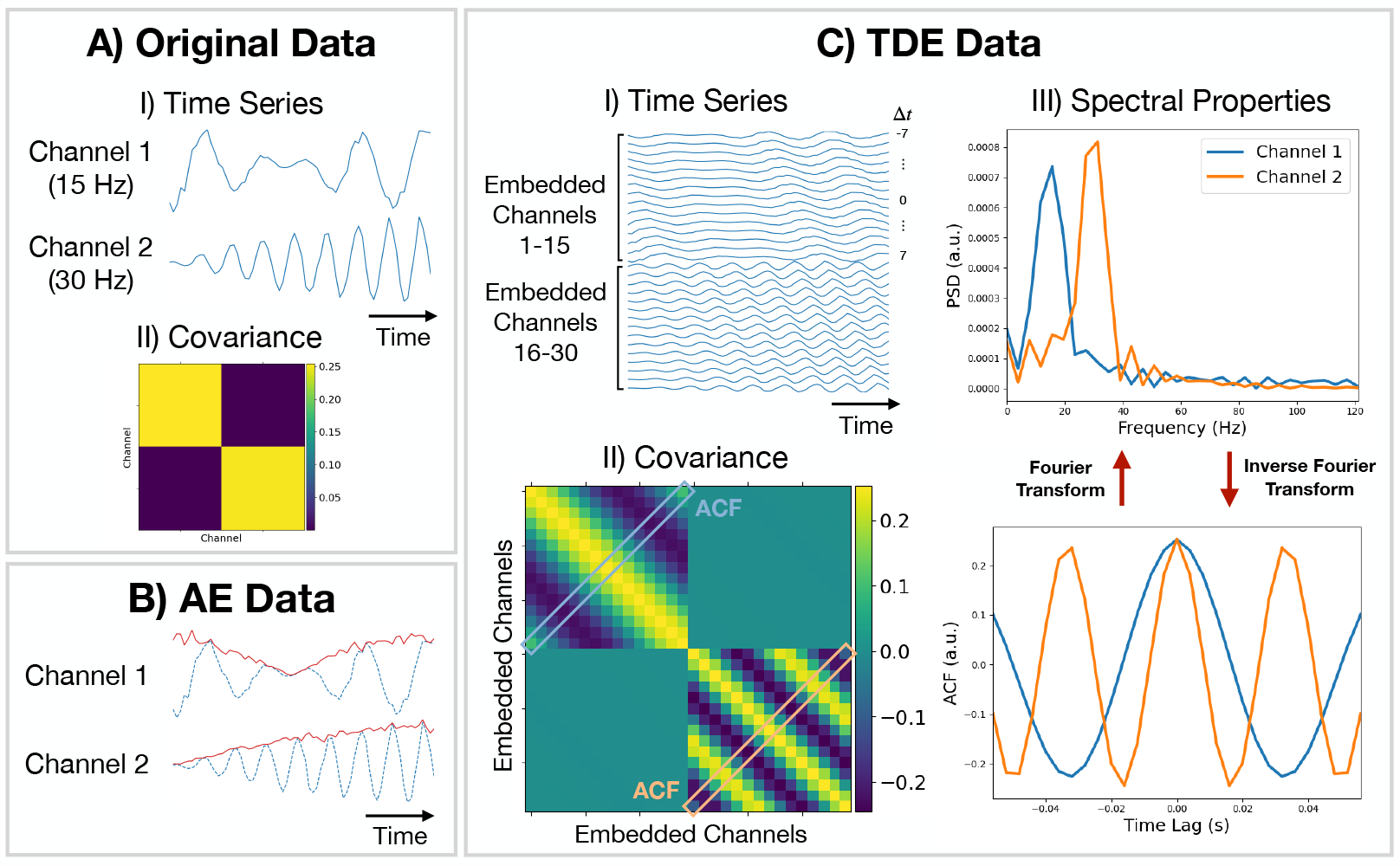
Methods for preparing training data. A.I) Original (simulated) time series data. Only a short segment (0.2 s) is shown. Channel 1 (2) is a modulated sine wave at 15 Hz (30 Hz) with *N* (0, 0.1) noise added. A.II) Covariance of the original data. B) Amplitude Envelope (AE) data (solid red line) and original data (dashed blue line). C.I) Time-Delay Embedded (TDE) time series. An embedding window of *±*7 lags was used. C.II) Covariance of TDE data. C.III) Spectral properties of the original data estimated using the covariance matrix of TDE data. Acronyms: Autocorrelation Function (ACF), Power Spectral Density (PSD).

**Time-Delay Embedding (TDE)** Studying the amplitude dynamics of oscillations does not reveal any insights into how different regions interact via phase synchronisation. For this, we need to prepare the data using TDE [43]. This augments the time series with extra channels containing time-lagged versions of the original channels. Figure 3C.I shows an example of this. To perform TDE, we need to specify the number of lagged channels to add (number of embeddings) and the lag to shift each additional channel by. In `osl-dynamics`, we always shift by one time point, so we only need to specify the number of lags. By adding extra channels, we embed the autocorrelation function (ACF) of the original data (as well as the cross-correlation function) into the covariance matrix of the TDE data. This is illustrated in Figure 3C.II. We plot the ACF taken from the TDE covariance matrix and the PSD (calculated using a Fourier transform) in Figure 3C.III. By using TDE data we make the covariance matrix sensitive to the frequency of oscillations in the original data. The covariance matrix is also sensitive to cross channel phase synchronisation via of the off-diagonal elements. Training on TDE data allows us to study dynamics in oscillatory amplitude and phase synchronisation between channels. When we prepare TDE data, we are normally only interested in looking for dynamics in the auto/cross correlation function via the covariance matrix, so we fix the mean to zero in the generative model.

For further details and example code for preparing data in `osl-dynamics` see the tutorial: https://osl-dynamics.readthedocs.io/en/latest/tutorials_build/data_preparation.html.

### 2.5 First-Level and Group-Level Analysis

Starting from the source reconstructed data, we study a dataset with a two-stage process:

1. **First-level analysis**. Here, our objective is to estimate subject-specific quantities. In the static (time-averaged) analysis, we calculate these quantities directly from the source data. However, if we are doing a dynamic analysis, we first train a generative model, such as the HMM or DyNeMo. Note, the HMM/DyNeMo are typically trained as group-level models using the concatenated data from all subjects. This means the models are unaware that the data originates from different subjects and allows the model to pool information across subjects, which can lead to more robust estimates of dynamic quantities. We use the latent description provided by the model with the source data to estimate the quantities of interest - this approach is known as *dual estimation*^7^ [44].
2. **Group-level analysis**. Quantities estimated for individual subjects, such as network metrics or summary statistics for dynamics, are used to model a group. For example, this could be predicting behavioural traits or characteristics of individual subjects, comparing two groups, or calculating the group average of an network response to a task. Typically, statistical significance testing is done at the group-level to verify that any observed differences or inferred relationships are not simply due to chance.

We will present the results of applying five pipelines to source reconstructed data calculated from the datasets mentioned in Section 2.2: a burst detection pipeline based on the HMM (discussed in Section 2.5.1); three dynamic network analysis pipelines based on the HMM and DyNeMo (discussed in Section 2.5.2) and a static network analysis pipeline (discussed in Section 2.5.3). Both the HMM and DyNeMo have been validated on simulated data for each application. See [45] for a demonstration of the HMM’s ability to identify oscillatory bursts and [24, 26] for a demonstration of the HMM/DyNeMo’s ability to identify dynamic networks.

#### 2.5.1 Burst Detection

We use an approach based on the HMM to detect bursts of oscillatory activity. In this approach, we prepare the source data using TDE. A typical TDE-HMM burst detection pipeline is shown in Figure 4. When the HMM state time courses are inferred on the training data, each “visit” to a particular state corresponds to a burst, or transient spectral event, with spectral properties specific to the state (e.g. an increase in *β*-band power). This approach assumes that we are looking for bursting in a single channel (brain region) at a time; separate HMMs can be used to detect bursting in each channel. We use the state time course to calculate summary statistics that characterise the dynamics of bursts. Typical summary statistics are:

- **Mean lifetime**^8^. This is the average duration a state is active.
- **Mean interval**. This is the average duration between successive state activations.
- **Burst count**. This is the number of times a state activates in a second on average.
- **Mean amplitude**. This is the average value of the AE of the source data when each state is active.

**Figure 4:**
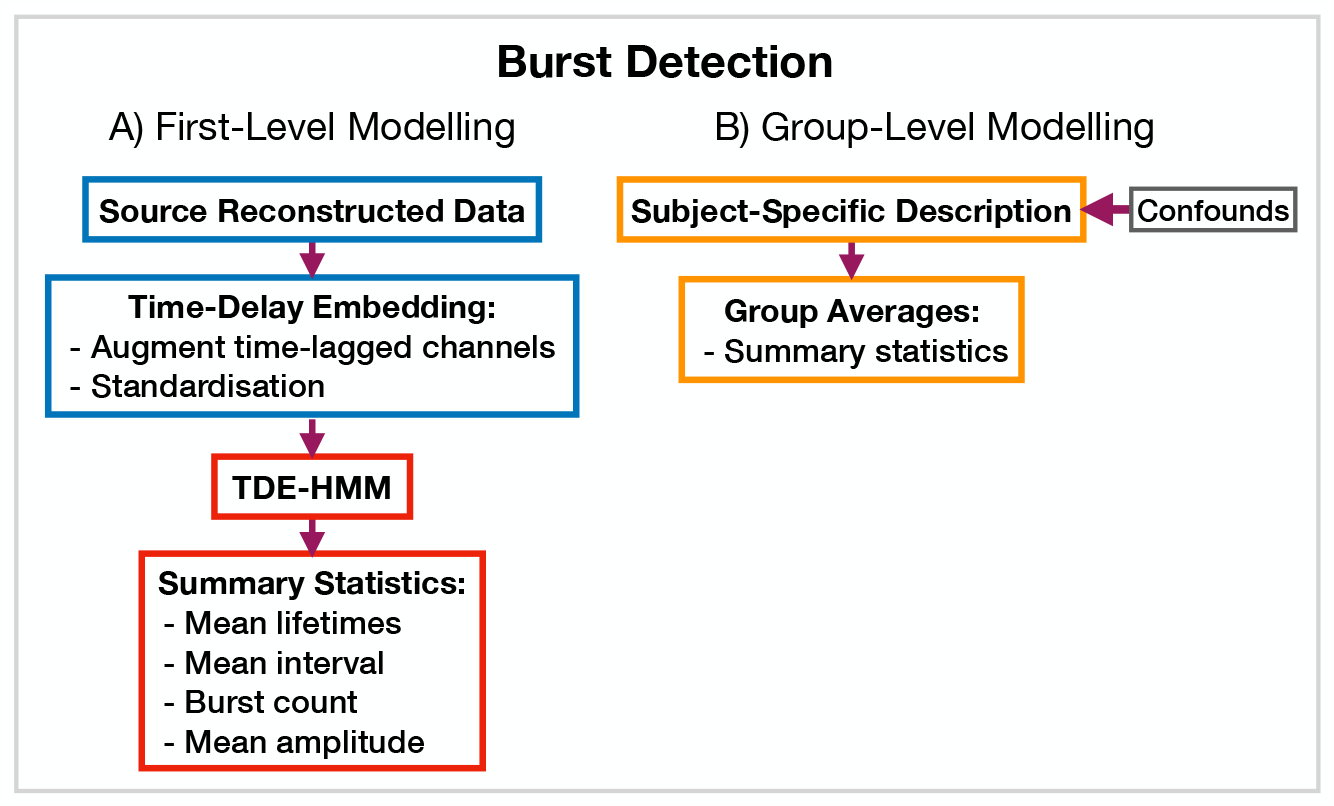
TDE-HMM burst detection pipeline. This is run on a single region’s parcel time course. Separate HMMs are trained for each region. A) Source reconstructed data is prepared by performing time-delay embedding and standardisation (z-transform). Following this an HMM is trained on the data and statistics that summarise the bursts are calculated from the inferred state time course. B) Subject-specific metrics summarising the bursts at a particular region are used in group-level analysis.

We calculate each of these for a particular state and subject. The averages are taken over all state activations. Given when a state is active we can use the source data to calculate the PSD of each burst type. We use the multitaper approach described in [24] to do this due to its ability to accurately estimate spectra. We present the results of applying a TDE-HMM burst detection pipeline to the CTF rest MEG dataset in Section 3.1.

#### 2.5.2 Identifying Dynamic Functional Networks

`osl-dynamics` provides more options for modelling dynamic functional networks. Note, in this case we train on multivariate data containing the activity at multiple regions of interest, rather than a single region, which is what we did in the burst detection pipeline (Section 2.5.1). Indeed, one perspective on using `osl-dynamics` to model dynamic functional networks, is that it is identifying bursts that span across multiple brain regions. Figure 5 shows the different combinations of data preparation and generative models that are available for a dynamic network analysis pipeline. We discuss each of these options and when they should be used below.

**Figure 5:**
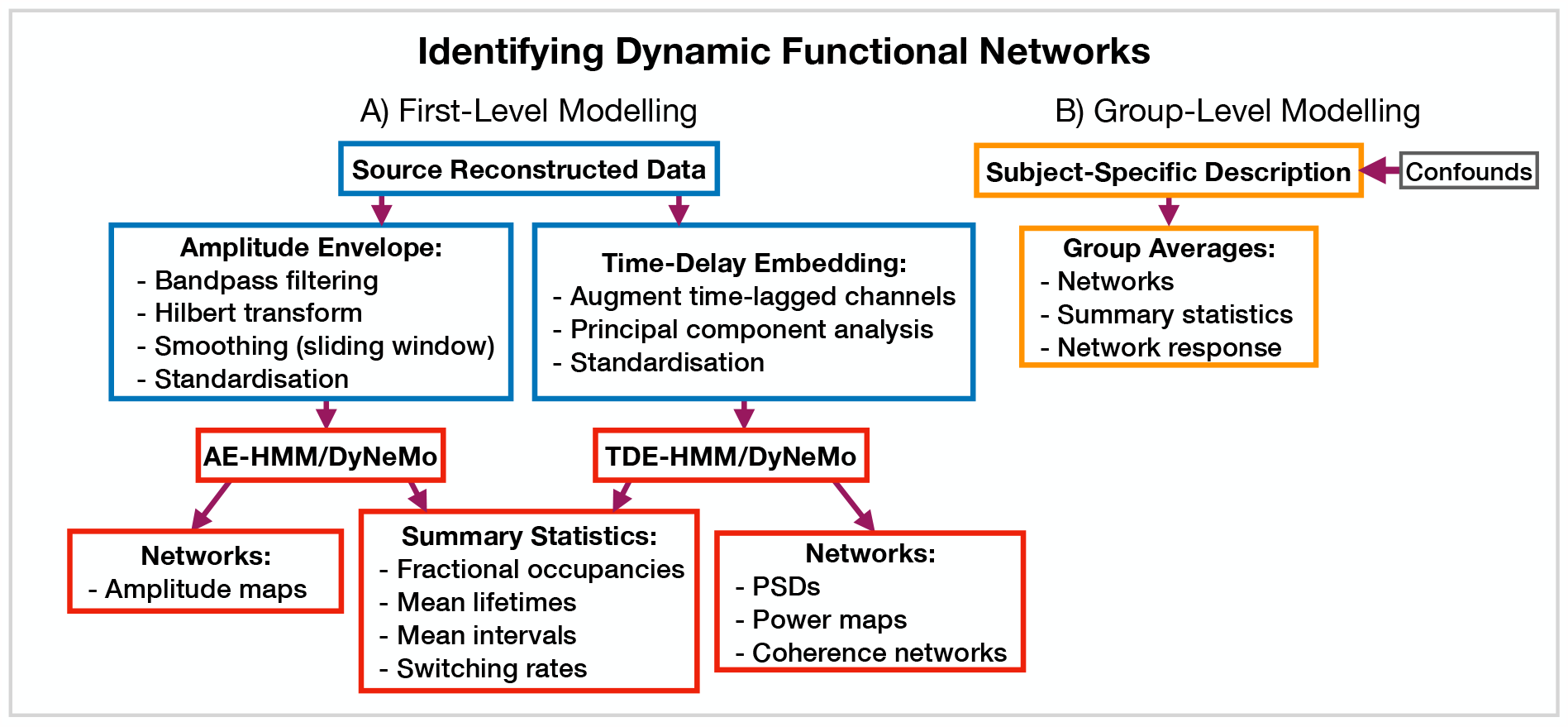
Dynamic functional network analysis pipeline. A) First-level modelling. This includes data preparation (shown in the blue boxes), model training and post-hoc analysis (shown in the red boxes). The first-level modelling is used to derive subject-specific quantities. B) Group-level modelling. This involves using the subject-specific description from the first-level modelling to model a group.

**AE-HMM**. If we are interested in identifying dynamics in amplitude, we can train on AE data. Once we have trained a model, we can estimate subject and state-specific networks (amplitude maps) using the training data and inferred state time course. Additionally, we can calculate summary statistics that characterise the dynamics from the inferred state time course. These summary statistics are:

- **Fractional occupancy**. This is the fraction of the total time that each state is active.
- **Mean lifetime**. This is the average duration that a state is active.
- **Mean interval**. This is the average duration between successive state visits.
- **Switching rate**. This is the number of activations per second of a particular state.

We calculate each of these for a particular state and subject. The averages are taken over all state activations. We present the results of an AE-HMM pipeline on the Elekta task MEG dataset in Section 3.2.

**TDE-HMM**. We can use TDE data to study dynamics in phase synchronisation as well as dynamics in amplitude. In a dynamic network analysis pipeline we train on a multivariate time series (i.e. the time series for all regions of interest together). This means after TDE we have a very large number of channels (number of embeddings times number of regions). Therefore, we often need to perform principal component analysis (PCA) for dimensionality reduction to ensure the data fits into computer memory.

In the TDE-HMM pipeline, we can calculate the same summary statistics as the AE-HMM pipeline. However, to estimate the functional networks we use the multitaper approach described in [24]. Here, we use the source data and inferred state time course to estimate subject, region and state-specific PSDs and cross PSDs. When then use the PSDs to calculate power maps and cross PSDs to calculate coherence networks, see [24] for further details. Note, we also use the spectral decomposition approached introduced in [14] to specify a frequency range for calculating power maps and coherence networks. This involves applying non-negative matrix factorisation to the stacked subject and state-specific coherence spectra to identify common frequency bands of coherent activity. In this report, we fit two spectral components and only present the networks for the first band, which typically cover 1-25 Hz. We will see the results of applying a TDE-HMM pipeline for dynamic network analysis on both the CTF rest and Elekta task MEG dataset in Section 3.3.

**TDE-DyNeMo**. This this pipeline, we replace the HMM with DyNeMo and train on TDE data. Unlike the mutually exclusive state description provided by the HMM, DyNeMo infers mode time courses, which describe the mixing ratio of each mode at each time point [26]. This mixture description complicates the calculation of subject-specific quantities, such as networks and summary statistics. To calculate mode and region-specific PSDs, we use the approach based on the General Linear Model (GLM) proposed in [46] where we regress the mixing coefficients onto a (cross) spectrogram, see [26] for further details. We then use the mode PSDs and cross PSDs to calculate power maps and coherence networks respectively. We can summarise the dynamics of each mode time course with quantities such as the mean, standard deviation and pairwise Pearson correlation. Alternatively, if we were interested in calculating the same summary statistics as the HMM (fractional occupancy, lifetime, interval, switching rate) we would first need to binarise the mode time courses. This can be done using a two-component Gaussian Mixture Model (GMM), which is discussed in [26]. Note, an additional complication related to the mode time course is that it does not contain any information regarding the relative magnitude of each mode covariance. For example, a mode with a small value for the mixing ratio can still be a large contributor to the instantaneous covariance if the values in the mode covariance matrix are relatively large. We account for this by renormalising the mode time course^9^, this is discussed further in [26]. We present the results of a TDE-DyNeMo pipeline on the CTF rest MEG dataset in Section 3.4.

AE-DyNeMo. The final option is to train DyNeMo on AE data. In this case, the amplitude maps are calculated using the GLM approach by regressing the mixing coefficients on a sliding window AE time course. Summary statistics for dynamics are calculated in the same way as the TDE-DyNeMo pipeline.

When we display the networks inferred by each of the pipelines above, we will threshold them to only show the strongest connections. In this work, we will specify the threshold using a datadriven approach where we fit a two-component GMM to the distribution of connections in each network. We interpret one of the components as the distribution for background connections and the other as the distribution for atypically strong connections, which is what we display in each plot.

#### 2.5.3 Identifying Static Functional Networks

A feature of `osl-dynamics` is that more conventional, static (time-averaged), network analyses can be carried out using the same methodology that we use in the dynamic methods. This allows for a much more straightforward comparison between static and dynamic analyses. To model static functional networks we simply need to specific the metrics we would like to use to summarise the networks and we calculate these directly from the source data. Figure 6 shows a typical static network analysis pipeline. We present the result of a static network analysis pipeline on the CTF rest MEG dataset in Section 3.5. Note, for the static networks we select the top 5% of connections to display in each plot rather than the GMM approached we used to threshold the dynamic functional networks.

**Figure 6:**
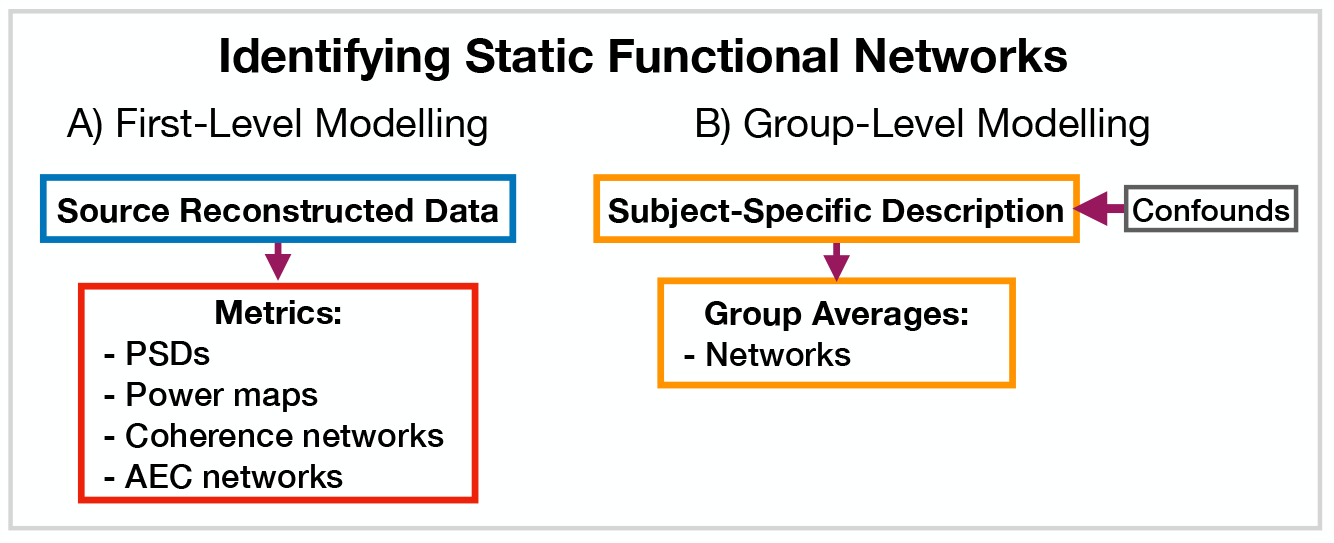
Static functional network analysis pipeline. A) The source reconstructed data is used to calculate metrics that describe networks. B) The subject-specific metrics are used to model a group. Acronyms: amplitude envelope correlation (AEC), power spectral density (PSD).

### 2.6 Run-to-Run Variability

The HMM and DyNeMo are trained by minimising a cost function (in `osl-dynamics`, we use the *variational free energy* [34, 26]). As is typical, this approach suffers from a local optimum issue, where the model can converge to different explanations (latent descriptions) of the data during training. I.e., different state/mode time courses can lead to similar values for the variational free energy. The final description can be sensitive to the stochasticity in updating model parameters and the initial parameter values.

A strategy for dealing with this that has worked well in the past is to train multiple models from scratch (each model is referred to as a *run*) and only the model with the lowest variational free energy is analysed. We consider this model as the best description of the data. We ensure any conclusions based on this model are reproducible in the best model from another set of independent runs. In all of our figures here, we present the results from the best run from a set of 10. In the supplementary information (SI) we show these results are reproducible in [47] an independent set of runs. Other strategies for dealing with run-to-run variability involve combining multiple runs, see for a discussion of these techniques.

## 3 Exemplary Analyses

In this section, we outline example uses of `osl-dynamics` to study source reconstructed MEG data. Section 3.1 presents the results of an oscillatory burst analysis pipeline. Sections 3.2-3.4 present the results of various dynamic network analysis pipelines. For comparison, we also include the results of a static network analysis pipeline in Section 3.5.

### 3.1 Burst detection using a single-region TDE-HMM

The pipeline in Figure 4 was applied to do burst detection on a single parcel in the left motor cortex. The source data was calculated using the CTF rest MEG dataset. All subjects were concatenated temporally and used to train the TDE-HMM. The results are shown in Figure 7. We see from the wavelet transform in Figure 7A.I that there are short bursts of oscillatory activity in this time series. This illustrates how it would be non-trivial, using conventional bandpass filtering and thresholding methods, to identify when exactly a burst occurs and what frequencies are contained within it. Instead of a conventional burst detection method, we use a 3 state TDE-HMM to identify bursts in a data-driven fashion. We see from the inferred state probability time course (Figure 7B.I) that there are short-lived states that describe this data. We can see from Figure 7B.II that each state corresponds to unique oscillatory activity. State 1 is interpreted as a non-oscillatory background state because it does not show any significant peaks in its PSD. States 2 and 3 show oscillatory activity in the 5/*θ* band (1-7 Hz) and α/, β band (7-30 Hz) respectively. Figure 7B.III shows the correlation of each state probability time course with the AEs for different frequency bands (Figure 7A.II). Based on this, we identify state 2 as a 5/*θ*-burst state and state 3 as a, β-burst state. We can see from Figure 7B.IV that these bursts have a variety of lifetimes ranging from a hundred to several hundred milliseconds. The reproducibility of these results is shown in SI Figure S1.

**Figure 7:**
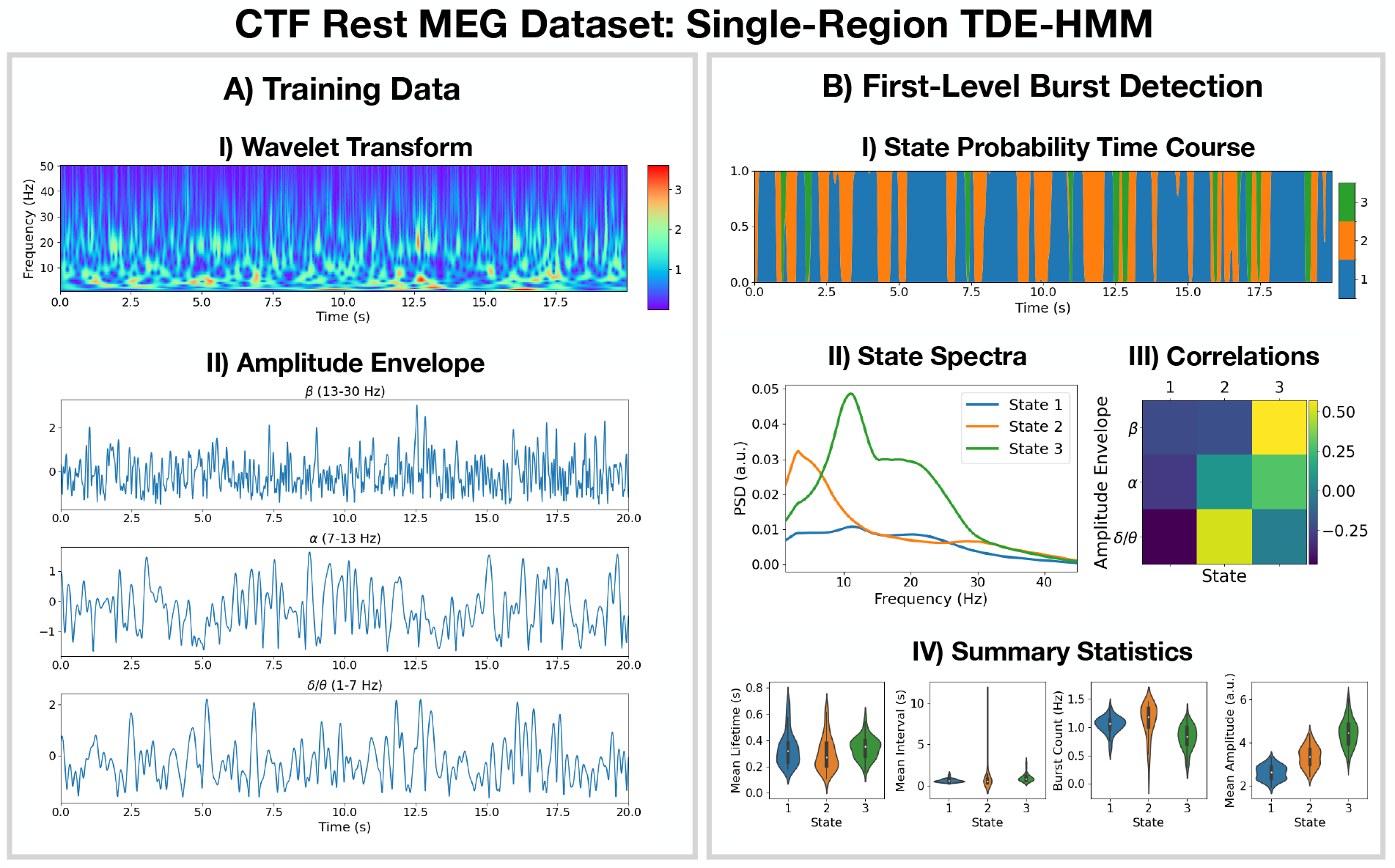
Burst detection: single region source reconstructed MEG data (left motor cortex) shows short-lived bursts of oscillatory activity. A.I) Dynamic spectral properties of the first 20 s of the time series from the first subject. A.II) Amplitude envelope calculated after bandpass filtering the time series over the,β-band (top), α-band (middle) and 5/*θ*-band (bottom). B.I) The inferred state probability time course for the first 20 s of the first subject. B.II) The PSD of each state. B.III) Pearson correlation of each state probability time course with the amplitude envelopes for different frequency bands. B.IV) Distribution over subjects for summary statistics characterising the bursts. Note, no additional bandpass filtering was done to the source data when calculating the mean amplitude. The script used to generate the results in this figure is here: https://github.com/OHBA-analysis/osl-dynamics/blob/main/examples/toolbox_paper/ctf_rest/tde_hmm_bursts.py.

### 3.2 Detecting network dynamics using a multi-region AE-HMM

The AE-HMM pipeline in Figure 5 was applied to source reconstructed data from the Elekta task MEG dataset to identify amplitude-based network dynamics. All subjects and runs were concatenated temporally to train the model. The results are shown in Figure 8 with an example of a group-level analysis on the HMM state time courses (calculation of a group-averaged network response).

**Figure 8:**
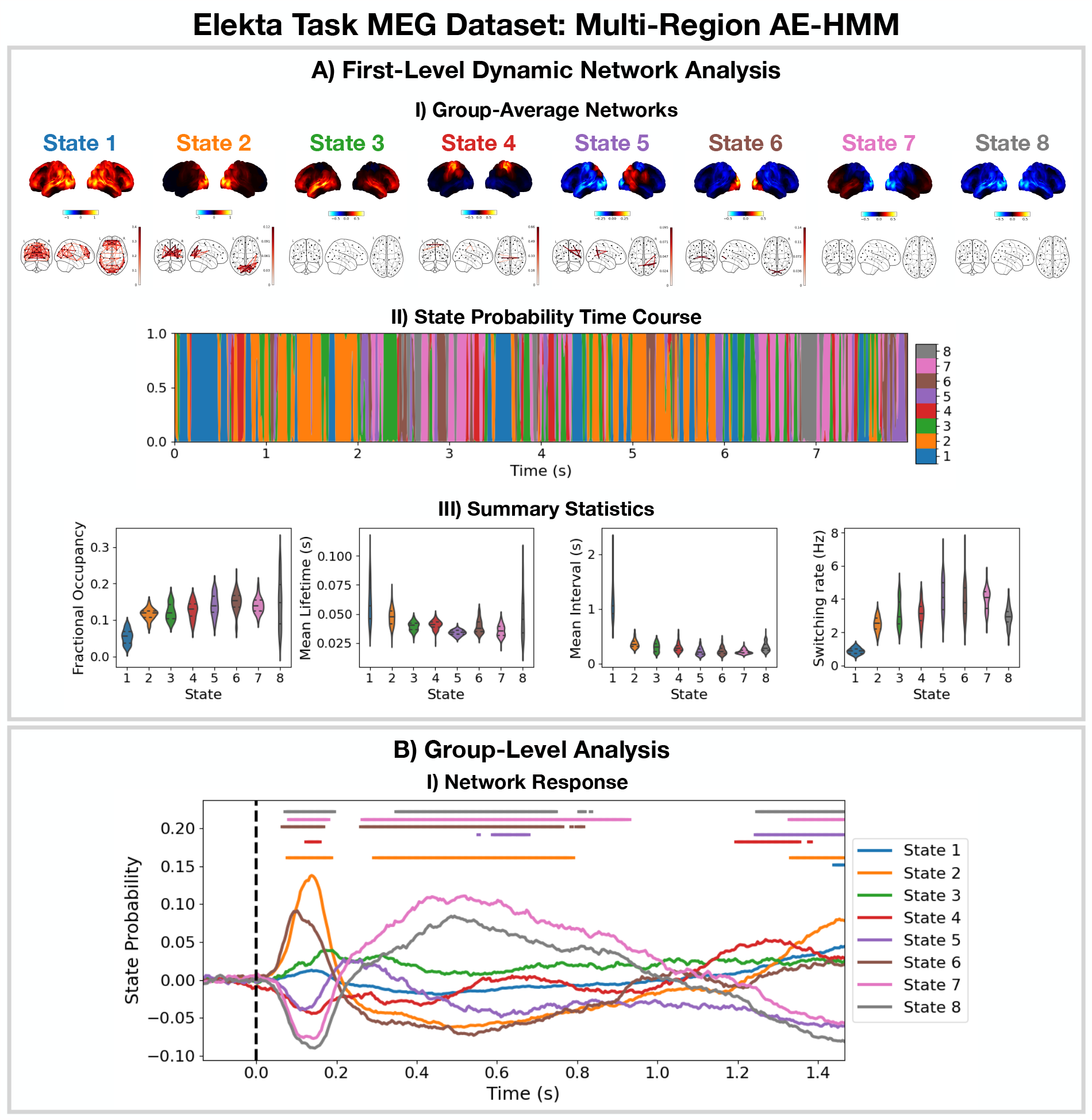
Dynamic network detection: a multi-region AE-HMM trained on the Elekta task MEG dataset reveals functional networks with fast dynamics that are related to the task. A.I) For each state, group-averaged amplitude maps relative to the mean across states (top) and absolute amplitude envelope correlation networks (bottom). A.II) State probability time course for the first 8 seconds of the first subject. A.III) Distribution over subjects for the summary statistics for each state. B.I) State time courses (Viterbi path) epoched around the presentation of visual stimuli. The horizontal bars indicate time points with p-value < 0.05. The maximum statistic pooling over states and time points was used in permutation testing to control for the family-wise error rate. The script used to generate the results in this figure is here: https://github.com/OHBA-analysis/osl-dynamics/blob/main/examples/toolbox_paper/elekta_task/tde_hmm.py.

We see the AE-HMM identifies plausible functional networks [48] with fast dynamics, typically with lifetimes of around 50 ms (Figure 8A.III). We identify a default mode network (state 1); two visual networks (states 2 and 6); a frontotemporal networks (state 3 and 7); and a sensorimotor network (state 4).

The AE-HMM was trained on the continuous source reconstructed data in an *unsupervised* manner, i.e. without any knowledge of the task. Post-HMM training, we can epoch the inferred state time course (Veterbi path) around the task (presentation of a visual stimuli^10^) and average over trials. This gives the probability of each state being activate around a visual event. This is shown in Figure 8B.I. We observe a significant increase (*p*-value < 0.05) in the activation of the visual networks (states 2 and 6) between 50-100 ms after the presentation of the as expected. We also observe a significant activation (*p*-value < 0.05) of the frontotemporal network (state 7) 300-900 ms after the visual stimuli as well as a deactivation of the visual networks (states 2 and 6). The reproducibility of these results is shown in SI Figure S2.

### 3.3 Detecting network dynamics using a multi-region TDE-HMM

The TDE-HMM pipeline in Figure 5 was also applied to the Elekta task MEG dataset. All subjects and runs were concatenated temporally and used to train the model. The results are shown in Figure 9.

**Figure 9:**
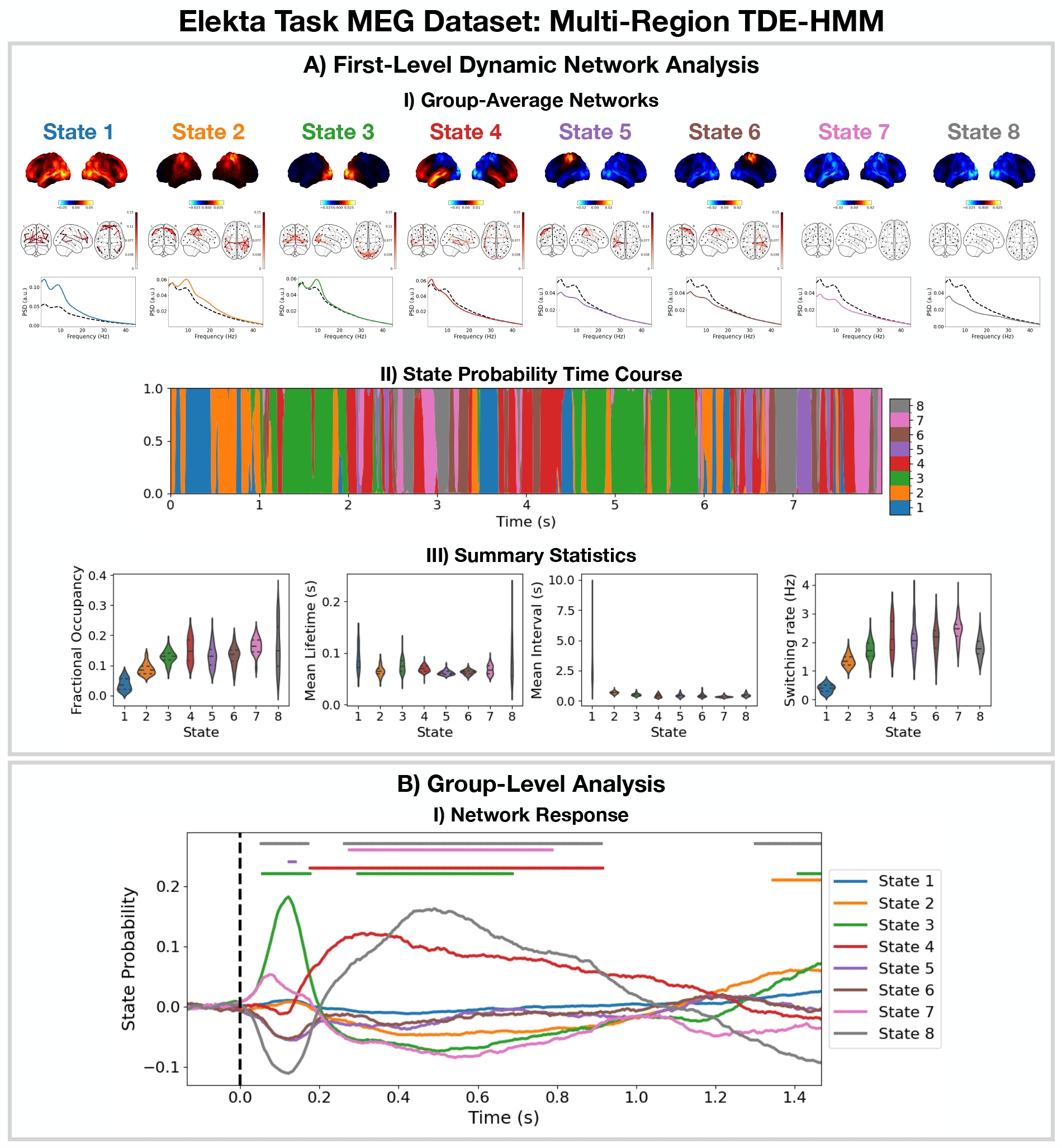
Dynamic network detection: a multi-region TDE-HMM trained on the Elekta task MEG dataset reveals spectrally distinct functional networks with fast dynamics. A.I) For each state, group-averaged power maps relative to the mean across states (top), coherence networks (middle) and PSD averaged over regions (bottom), both the statespecific (coloured solid line) and static PSD (i.e. the average across states, dashed black line) are shown. A.II) State probability time course for the first 8 seconds of the first subject. A.III) Distribution over subjects for the summary statistics for each state. B.I) State time courses (Viterbi path) epoched around the presentation of visual stimuli. The horizontal bars indicate time points with p-value < 0.05. The maximum statistic pooling over states and time points was used in permutation testing to control for the family-wise error rate. The script used to generate the results in this figure is here: https://github.com/OHBA-analysis/osl-dynamics/blob/main/examples/toolbox_paper/elekta_task/tde_hmm.py.

For the high-power networks (states 1-4), we see the same spatial patterns in TDE-HMM power maps (Figure 9A.I, top) and AE-HMM amplitude maps (Figure 8A.I). We can see from the state PSDs (Figure 9A.I, bottom) that the networks identified by the TDE-HMM exhibit distinct spectral (oscillatory) activity. The TDE-HMM networks also have fast dynamics (Figure 9A.III) with lifetimes of around 50 ms. In Figure 9B.I, we can see we are able to reproduce the network response analysis we did using the AE-HMM (Figure 8B.I). The reproducibility of these results is shown in SI Figure S3.

The Elekta MEG dataset was recorded during a visual perception task. For comparison, we perform the same analysis on the CTF rest MEG dataset. All subjects were concatenated temporally and used to train the model. Figure 10 shows the results of applying a TDE-HMM pipeline to this dataset. We observe similar networks in rest (Figure 10A) as in task (Figure 9A), which is a known result from fMRI studies [49]. We also include the coherence networks in Figures 9A.I and 10A.I. We observe regions with high power activations have high connectivity (coherence). These networks also have fast dynamics (Figure 10A.III) with lifetimes of 50-100 ms.

**Figure 10:**
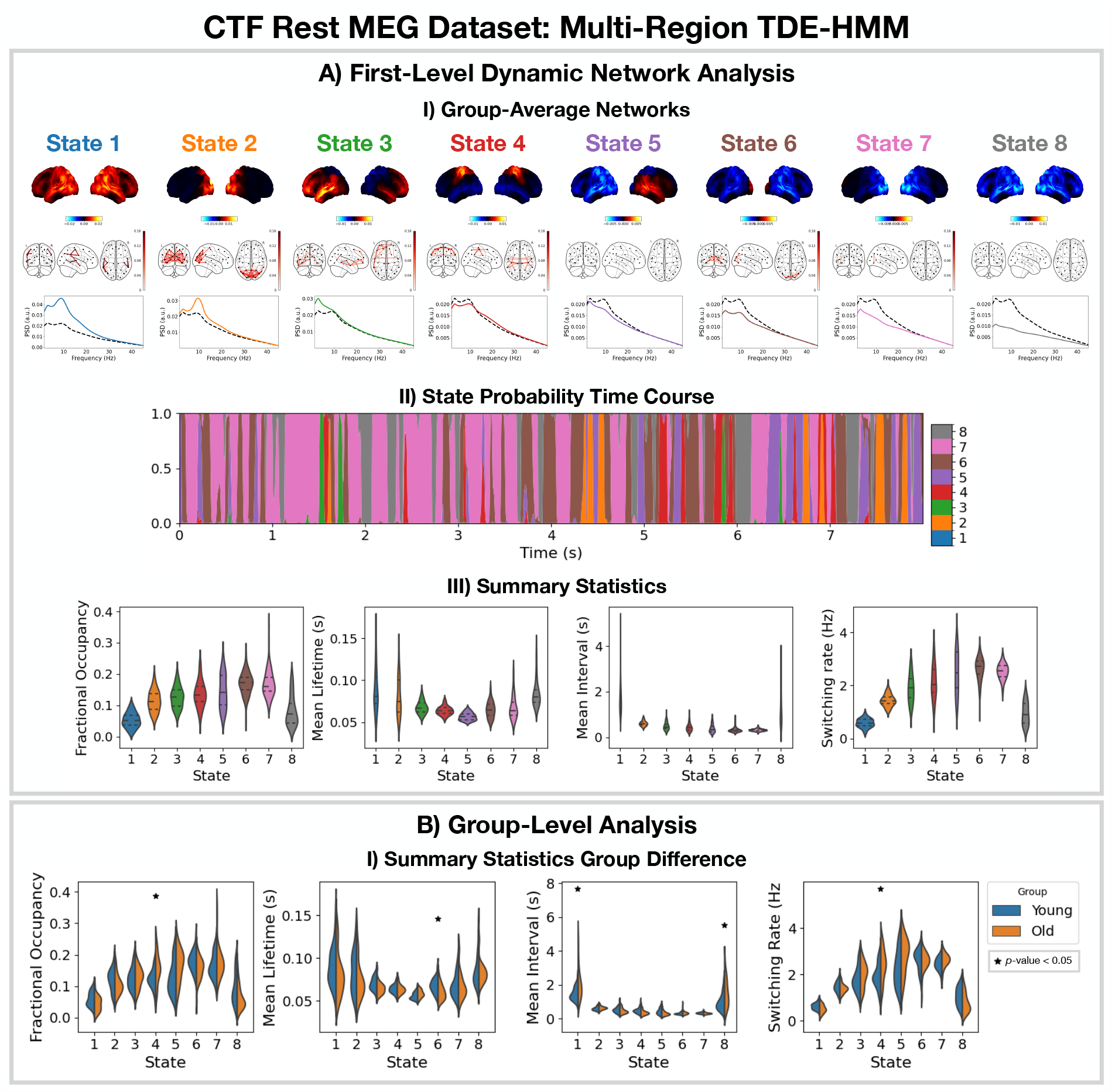
Dynamic network detection: a multi-region TDE-HMM trained on the CTF rest MEG dataset identifies the same functional networks to those found with the Elekta task MEG dataset and reveals differences in the dynamics for young vs old groups. A.I) For each state, group-averaged power maps relative to mean across states (top), absolute coherence networks (middle) and PSD averaged over regions (bottom), both the state-specific (coloured solid line) and static PSD (i.e. the average across states, dashed black line) are shown. A.II) State probability time course for the first 8 seconds of the first subject and run. A.III) Distribution over subjects for the summary statistics of each state. B.I) Comparison of the summary statistics for a young (18-34 years old) and old (34-60 years old) group. The star indicates a p-value < 0.05. The maximum statistic pooling over states and metrics was used in permutation testing to control for the family-wise error rate. The script used to generate the results in this figure is here: https://github.com/OHBA-analysis/osl-dynamics/blob/main/examples/toolbox_paper/ctf_rest/tde_hmm_networks.py.

To illustrate a group-level analysis we could do with a dynamic network perspective, we compared two groups: 27 subjects in a young group (18-34 years old) and 38 subjects in an old group (34-60 years). Figure 10B.I shows summary statistics for each group. We see the fractional occupancy and switching rate of the sensorimotor network (state 4) is increased in the older group (*p*-value < 0.05). The mean lifetime of the visual network (state 6) is also decreased in the older group (p-value < 0.05). The older group also has a wider distribution of mean intervals for the default mode network (state 1) and suppressed state (8) (p-value < 0.05). The reproducibility of these results is shown in SI Figure S4. The age-related differences we observe here are consistent with existing studies [50]. We will discuss the young vs old comparison further in Section 3.5.

### 3.4 Dynamic network detection using multi-region TDE-DyNeMo

The TDE-DyNeMo pipeline in Figure 5 was applied to the CTF rest MEG dataset. All subjects were concatenated temporally and used to train the model. The results are shown in Figure 11. Note, for DyNeMo we found that learning 7 modes (rather than 8) led to more reproducible results. Therefore, we present the 7 mode fit in Figure 11.

**Figure 11:**
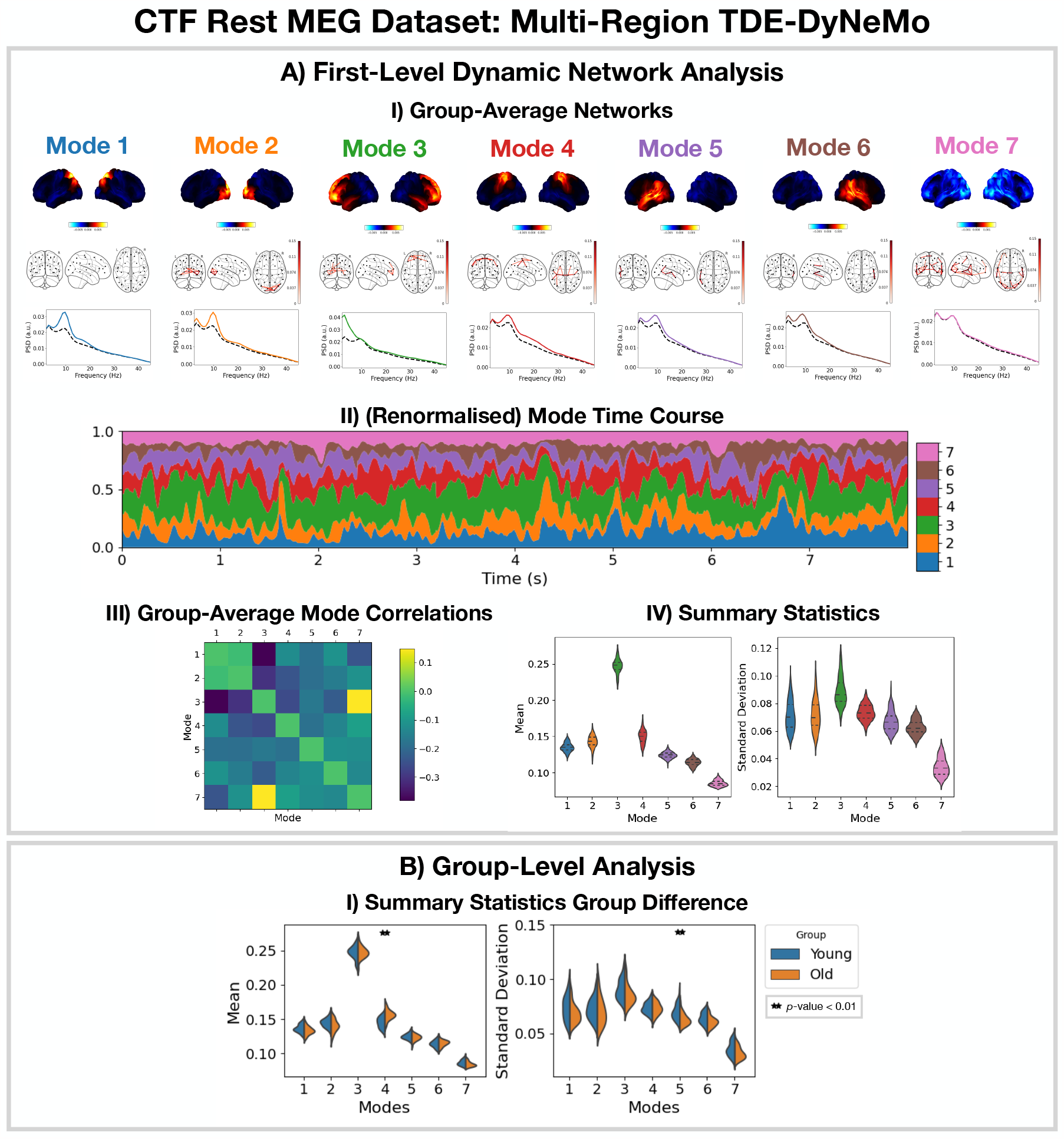
Dynamic network detection: a multi-region TDE-DyNeMo trained on the CTF rest MEG dataset reveals spectrally distinct modes that are more localised than HMM states and overlap in time. A.I) For each mode, group-averaged power maps relative to the mean across modes (top), absolute coherence networks (middle) and PSD averaged over regions (bottom), both the mode-specific (coloured solid line) and static PSD (i.e. the average across modes, dashed black line) are shown. A.II) Mode time course (mixing co-efficients) renormalised using the trace of the mode covariances. A.III) Pearson correlation between renormalised mode time courses calculated by concatenating the time series from eachsubject. A.IV) Distribution over subjects for summary statistics (mean and standard deviation) of the renormalised mode time courses. B.I) Comparison of the summary statistics for a young (18-34 years old) and old (34-60 years old) group. The maximum statistic pooling over modes and metrics was used in permutation testing to control for the family-wise error rate. The script used to generate the results in this figure is here: https://github.com/OHBA-analysis/osl-dynamics/blob/main/examples/toolbox_paper/ctf_rest/tde_dynemo_networks.py.

We can see from the power maps and coherence networks (Figure 11A.I) that DyNeMo identifies much more localised power activations and a cleaner network structure than was seen with the TDE-HMM. We can see from the PSDs (Figure 11A.I, bottom) that these networks also exhibit distinct spectral characteristics.

From the (renormalised) mode time course (Figure 11A.II) we see the description provided by DyNeMo is that of overlapping networks that dynamically vary in mixing ratios. This is a complementary perspective to the state description provided by the HMM. Co-activations of each mode can be understood by looking at the Pearson correlation between (renormalised) mode time courses (Figure 11A.III). We observe modes with activity in neighbouring regions show more co-activation. We summarise the (renormalised) mode time course using statistics (the mean and standard deviation) in Figure 11A.IV.

To compare DyNeMo to the HMM in a group-level analysis, we repeat the young vs old study using the DyNeMo-specific summary statistics (i.e. the mean and standard deviation of the renormalised mode time courses). Figure 11B.I shows significant group differences for young (18-34 years old) and old (34-60 years old) participants. We can see an increased mode contribution (mean renormalised mode time course) for the sensorimotor network (mode 4), which reflects the increase in fractional occupancy we saw in the TDE-HMM (Figure 10B.I). We see DyNeMo is able to reveal a stronger effect size with a *p*-value < 0.01 compared to the TDE-HMM, which had a *p*-value < 0.05. DyNeMo also shows a decrease in the variability (standard deviation of the renormalised mode time course) for the left temporal network (mode 5, *p*-value < 0.01). We will discuss the young vs old comparison further in Section 3.5. The reproducibility of these results is shown in SI Figure S5.

### 3.5 Estimating Static Functional Networks

For comparison, we also apply a typical static network analysis pipeline (including static functional connectivity) to the CTF rest MEG dataset. We also consider how the static perspective in a young vs old group-level analysis compares to the dynamic perspective provided by the TDE-HMM in Figure 10 and TDE-DyNeMo in Figure 11, illustrating the benefits of being able to do static and dynamic analyses within the same toolbox.

Figure 12 shows the the group-averaged PSD (A.I), power maps (A.II), coherence networks (A.III) and amplitude envelope correlation (AEC) networks (A.IV) calculated using all subjects. We observe 5-power is strongest in anterior regions and α-power is strongest in posterior regions. We also observe qualitatively similar coherence and AEC networks. In particular, we see strong occipital connectivity in the α-band in both the coherence and AEC networks.

**Figure 12:**
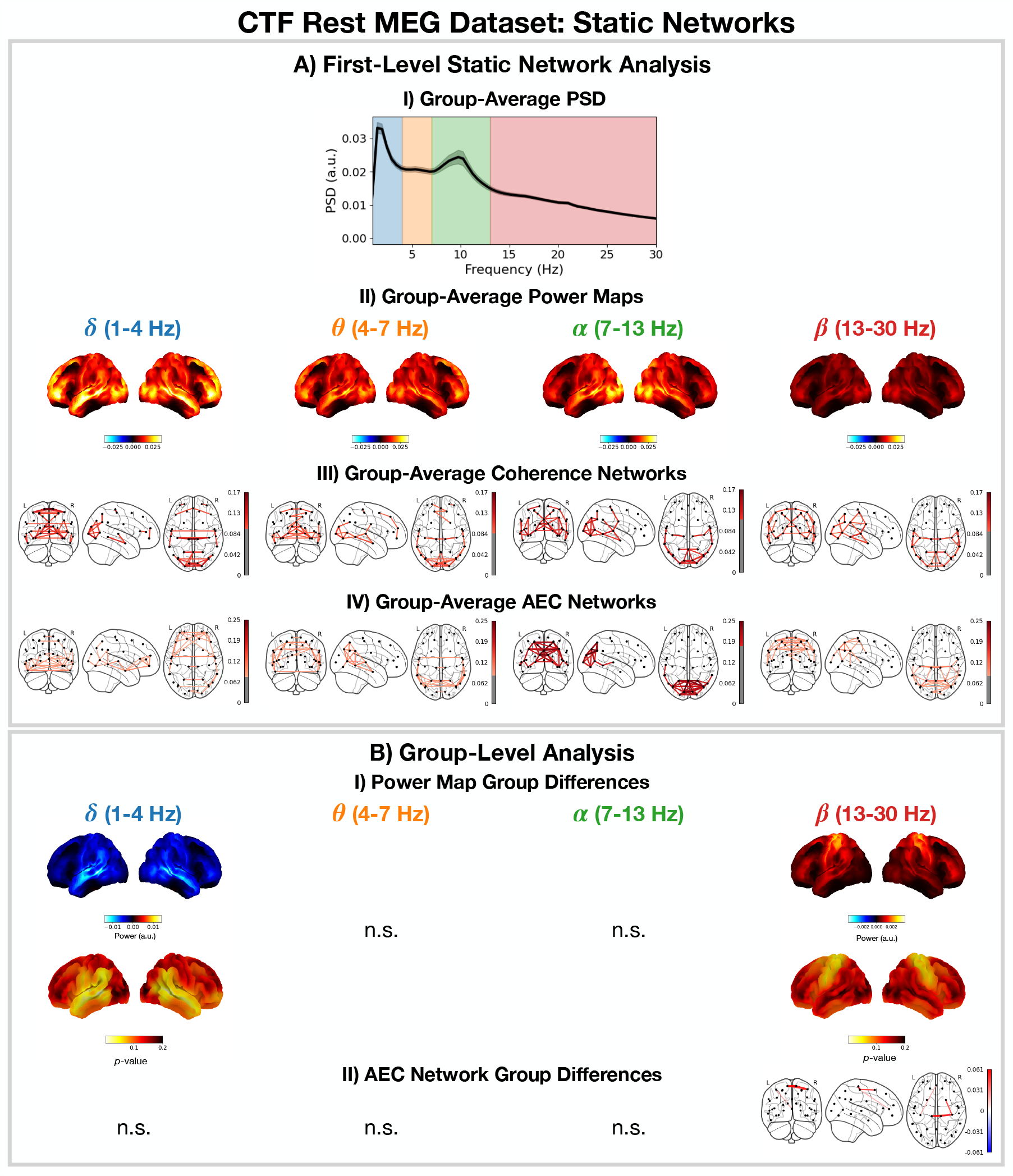
Static network detection: osl-dynamics can also be used to perform static network analyses (including functional connectivity). In the CTF rest MEG dataset, this reveals frequency-specific differences in the static functional networks of young (18-34 years old) and old (34-60 years old) participants. Group-average PSD (averaged over subjects and parcels, A.I) power maps (A.II) coherence networks (A.III) and AEC networks (A.IV) for the canonical frequency bands (*δ, θ, α, β*. B.I) Power difference for old minus young (top) and p-values (bottom). Only frequency bands with at least one parcel with a *p*-value < 0.05 are shown, the rest are marked with n.s. (none significant). B.II) AEC difference for old minus young only showing edges with a p-value < 0.05. The maximum statistic pooling over frequency bands and parcels/edges was used in permutation testing to control for the family-wise error rate. The script used to generate the results in this figure is here: https://github.com/OHBA-analysis/osl-dynamics/blob/main/examples/toolbox_paper/ctf_rest/static_networks.py.

Figure 12B shows significant (*p*-value < 0.05) differences in the power maps (B.I) and AEC networks (B.II) for old (34-60 years old) minus young (18-34 years old) groups. We observe a significant reduction in temporal *δ*-power and increase in sensorimotor *β*-power. We also observe a significant increase in sensorimotor AEC in the, *β*-band (Figure 12B.II).

The static (time-averaged) differences we see in young vs old participants can arise in many ways from the underlying dynamics of resting-state networks (Figure 10A.I and 11A.I). For example, an increase in static power could due to more frequent activations of a particular network. Conversely, the dynamics of the networks may be unaffected and the power within a network could be altered. Studying the dynamic network perspective using the HMM and/or DyNeMo can help provide further insights into how the static differences arise. Looking at the dynamic network perspective provided by the TDE-HMM, we see an increase in the fractional occupancy of state 4 (Figure 10B.I), which is a network with high *β*-power and connectivity (coherence) in the sensorimotor region. This is consistent with the static increase in *β*-power and AEC connectivity we observe here; i.e. the increase in static, *β*-power and connectivity with age can be linked to a larger fraction of time spent in the sensorimotor network. The perspective provided by TDE-DyNeMo shows an increase with age in the contribution of mode 4 (Figure 11B.I), which represents a sensorimotor network. This is a complementary explanation for the increase in static, *β*-power and connectivity as a larger contribution from the sensorimotor network to the overall brain activity of older participants.

## 4 Discussion

In Section 3.1, we use the TDE-HMM to identify oscillatory bursts in a data-driven manner with much fewer assumptions than conventional burst detection methods based on amplitude thresholding. The advantages of using a data-driven approach like the TDE-HMM are discussed further in [9, 51]. In short, with a conventional approach we must pre-specify a frequency of interest and we may miss oscillatory bursts that do not reach an arbitrary threshold. In contrast, the TDE-HMM is less sensitive to the amplitude (it is better able to identify low-amplitude oscillatory bursts) and can identify the frequency of oscillations automatically.

In Sections 3.2 and 3.3, we presented the functional networks identified by HMMs in a variety of settings. These networks were identified automatically at fast (sub-second) timescales from the data (unsupervised) with no input from the user. We found a set of plausible networks that were related to task (Figure 8) and demographics (Figure 10). These networks were very reproducible: across multiple HMM runs; across different data preparation techniques (AE and TDE); across different experimental paradigms (task and rest) and across different scanners (Elekta and CTF).

Given we observe similar networks with the AE-HMM and TDE-HMM (Figures 8 and 9 respectively), one may ask which pipeline is recommended. The TDE-HMM approach is able to model dynamics in oscillatory amplitude and phase synchronisation whereas the AE-HMM can only model dynamics in amplitude. This means the TDE-HMM is generally a better model for oscillatory dynamics. An occasion where the AE-HMM may be preferred is if the extra computational load of training on TDE/PCA data prohibits the TDE-HMM.

An important choice that has to be made when training an HMM/DyNeMo is the number of states/modes. For burst detection, we are often interested in identifying the time points when a burst occurs. This can be achieved by fitting a two state HMM: and ‘on’ and ‘off’ state. If we’re interested in multiple burst types, we can increase the number of states. In this work, we chose a three state HMM to stay close to the on/off description while allowing for multiple burst types. For the dynamic network analysis, we want a low number of states/modes to give us a compact representation of the data. A common choice is between 6 and 12. We can use the reproducibility analysis (section 2 in the SI) to show a given number of states/modes is reproducible and use this to find an upper limit for the number of states/modes that can be reliably inferred.

`osl-dynamics` offers a choice of two generative models for detecting network dynamics: the HMM and DyNeMo. The HMM assumes that there are mutually exclusive network states, whereas DyNeMo assumes the network modes are mixed differently at each time point. While DyNeMo’s assumption is arguably more realistic, the HMM’s stronger assumption has the benefit of simplifying the decomposition, which can make interpreting the network dynamics more straightforward. In short, the HMM and DyNeMo provide complementary descriptions of network dynamics, with either one being potentially useful depending on the context [26]. DyNeMo does have the additional advantage of using a richer temporal regularisation through the use of a deep recurrent network. This has been shown to capture longer range temporal structure than the HMM [26], and exploring the cognitive importance of long-range temporal structure is an interesting area of future investigation [52]. It is possible to quantify which model is better using a downstream task. In this manuscript, the downsteam tasks are the evoked network response and young vs old group differences. We argue a better performance in the downstream task indicates a more useful model.

`osl-dynamics` can also be used to compute static network descriptions, including conventional static functional connectivity. This uses the same methodology as the state (or mode) specific network estimation in the dynamic approaches, making comparisons between dynamic and static perspectives more straightforward. In Section 3.5, we used this feature to relate the static functional network description to a dynamic perspective. We would like to stress that the young vs old study is used as an example of the type of group analyses that can be performed with this toolbox and that a more rigorous study with a larger population dataset is needed to understand the impact of ageing on functional networks. The results in Section 3.5 should be taken as just an indication of possible ageing effects that can be investigated in a future study. In this report, we focus on the presentation of the tools needed to make such studies possible.

## 5 Conclusions

We present a new toolbox for studying time series data: `osl-dynamics`. This is an open-source package written in Python. We believe the availability of this package in Python improves the accessibility of these tools, in particular for non-technical users. Additionally, it avoids the need for a paid license. Using Python also enables us to take advantage of modern deep learning libraries (in particular `TensorFlow` [53]) which enables us to scale these methods to very large datasets, something that is currently not possible with existing toolboxes.

`osl-dynamics` can be used, and has been used, in a wide range of applications and on a variety of data modalities: electrophysiological, invasive local field potential, functional magnetic resonance imaging, etc. Here, we illustrated its use in applications of burst detection and dynamic network analysis using MEG data. This package also allows the user to study the static (time averaged) properties of a time series alongside dynamics within the same toolbox. The methods contained in `osl-dynamics` provide novel summary measures for dynamics and group-level analysis tools that can be used to inform our understanding of cognition, behaviour and disease.

## 6 Credit authorship contribution statement

CG: Conceptualisation, Methodology, Software, Validation, Formal analysis, Investigation, Data curation, Writing - original draft, Writing - review and editing, Visualisation. RH: Methodology, Software, Validation. ER: Software. MWJE: Methodology, Software. AJQ: Methodology. DV: Methodology, Writing - review and editing. MWW: Conceptualisation, Methodology, Data curation, Writing - review and editing, Supervision.

## 7 Acknowledgements

This research was supported by the National Institute for Health Research (NIHR) Oxford Health Biomedical Research Centre. The Wellcome Centre for Integrative Neuroimaging is supported by core funding from the Wellcome Trust (203139/Z/16/Z). CG is supported by the Wellcome Trust (215573/Z/19/Z). RH is supported by the EPSRC Centre for Doctoral Training in Health Data Science (EP/S02428X/1). ER is supported by an Engineering and Physical Sciences Research Council (EPSRC) and MRC grant (EP/L016044/1) and F. Hoffmann-La Roche. MWJE is supported by the Wellcome Trust (106183/Z/14/Z, 215573/Z/19/Z), the New Therapeutics in Alzheimer’s Diseases (NTAD) supported by the MRC and the Dementia Platform UK (RG94383/RG89702). DV is supported by a Novo Nordisk Foundation Emerging Investigator Fellowship (NNF19OC-0054895), an ERC Starting Grant (ERC-StG-2019-850404), and a DFF Project 1 from the Independent Research Fund of Denmark (2034-00054B). MWW is supported by the Wellcome Trust (106183/Z/14/Z, 215573/Z/19/Z), the New Therapeutics in Alzheimer’s Diseases (NTAD) study supported by UK MRC, the Dementia Platform UK (RG94383/RG89702) and the NIHR Oxford Health Biomedical Research Centre (NIHR203316). The views expressed are those of the author(s) and not necessarily those of the NIHR or the Department of Health and Social Care.

## Supplementary Information

### 1 Overview of osl-dynamics

In this section, we provide an overview of the `osl-dynamics` package. This include a description of the source code (Section 1.1), installation (Section 1.2), documentation and tutorials (Section 1.3).

Note, `osl-dynamics` utilises many popular open-source scientific computing and data science packages in python: `numpy, scipy, pandas, sklearn,` etc^11^. Virtually all the the plotting functions are wrappers for `matplotlib or nilearn. osl-dynamics` also uses the deep learning library `TensorFlow` [1] to implement the models. This package enables us to easily scale to large datasets and take advantage of graphical processing units (GPUs), which can speed up training a model by a factor of 10.

#### 1.1 Source Code

The source code is hosted on a public GitHub repository: https://github.com/OHBA-analysis/osl-dynamics. The different sub-packages and their use cases are:

- `analysis`: contains functions for post-hoc analysis; group-level modelling and statistical significance testing. This includes: functions for calculating spectral properties of time series data; fitting linear regressions; fitting Gaussian mixture models and General Linear Model (GLM) statistical significance testing using permutations.
- `config_api`: contains functions for using a simple user interface.
- `data`: useful classes and functions for managing and preparing neuroimaging data. This sub-package contains the Data class which is the central object used in osl-dyanmics for handling and preparing data.
- `files`: additional files for plotting and managing data, such as parcellation files and structural MRIs.
- `inference`: classes and functions that are helpful for training custom TensorFlow models. This includes custom classes for typical components in deep learning models, such as layers, initialisers, callbacks, etc.
- `models`: classes for available models. This include the HMM and DyNeMo, as well as other models that are under active development. Each model is implemented using keras [2] in TensorFlow.
- `simulation`: classes for simulating time series data with popular models. This includes classes for simulating HMMs and multivariate autoregressive models.
- `utils`: additional utility classes and functions. This sub-package contains the plotting module, which is helpful for visualising results.

#### 1.2 Installation

`osl-dynamics` can be installed on most modern computers in a similar fashion to other popular python packages via `pip` [3]. Installation instructions can be found here: https://osl-dynamics.readthedocs.io/en/latest/install.html. In short, `osl-dynamics` can be installed in three steps:

1. Create a virtual environment.
2. Install `TensorFlow` (and add ons).
3. Install `osl-dynamics`.

Note, the installation of `TensorFlow` is separate to `osl-dynamics` because there maybe computer/hardware specific differences which may require different installations of `TensorFlow`, see the `TensorFlow` documentation for further details [1].

For people who wish to contribute to development or would like to have access to the latest development version, `osl-dynamics` can also be installed from source using the instructions on the GitHub repository `README.md`.

### 1 Documentation and Tutorials

The official documentation for the `osl-dynamics` package is here: https://osl-dynamics.readthedocs.io/en/latest. The documentation contains a short description of the models contained in osl-dynamics, a page for frequently asked questions (FAQs) and an API reference guide, which lists all the modules/classes and functions in the package. The API reference guide is useful for looking up the inputs and outputs of various functions/classes. The documentation also includes comprehensive tutorials describing each pipeline (as well as other analyses such as sliding windows and estimation of static networks) in more detail.

### 2 Reproducibility

We use variational Bayesian inference to learn model parameters (state/mode time courses and observation model means/covariances). This process involves updating the model parameters to minimise the *variational free energy* [4].

Each time we train an HMM (Hidden Markov Model) or DyNeMo (Dynamic Network Modes), we start from random model parameters and use a stochastic procedure to update the parameters. This leads to some variability in the final model parameters we converge to and their corresponding variational free energy. We deem the model parameters with the lowest free energy as the ones that best describe the data and use these for subsequence analysis. We deem a set of results as reproducible if we are consistently able to infer the same model parameters from a dataset. Empirically, we find picking the best run from a set of 10 consistently finds the same set of model parameters. Note, for a given set of hyperparameters, only the relative difference between values for the variational free energy is important.

We show the variational free energy for 3 sets of 10 runs in Figures S1-S5 (A). In red we highlight the variational free energy for the best run (i.e. the one with the lowest value). Within a set of runs, we see large variability in the variational free energy which would reflect variability in the model parameters. Only looking at the best run across sets we see similar values for the variational free energy. In Figures S1-S5 (B), we see the summary statistics for the best run from each set is very similar indicating we infer the same dynamics in each best run. In Figures S2-S5 (C), we see we infer the same amplitude/power maps. In Figures S2-S5 (D), we see we observe the same results from the group-level analysis across all 3 sets.

**Figure S1:**
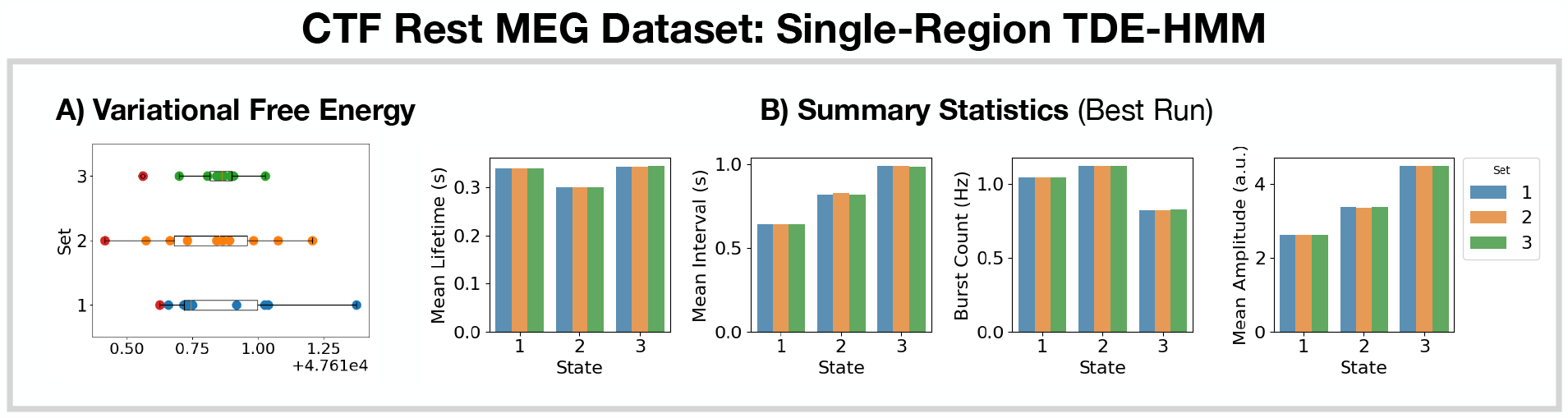
Reproducibility of the CTF rest MEG dataset single-region TDE-HMM analysis. A) Variational free energy for 3 sets of 10 runs. B) Summary statistics for the best run from each set (indicated by the red dot in A).

**Figure S2:**
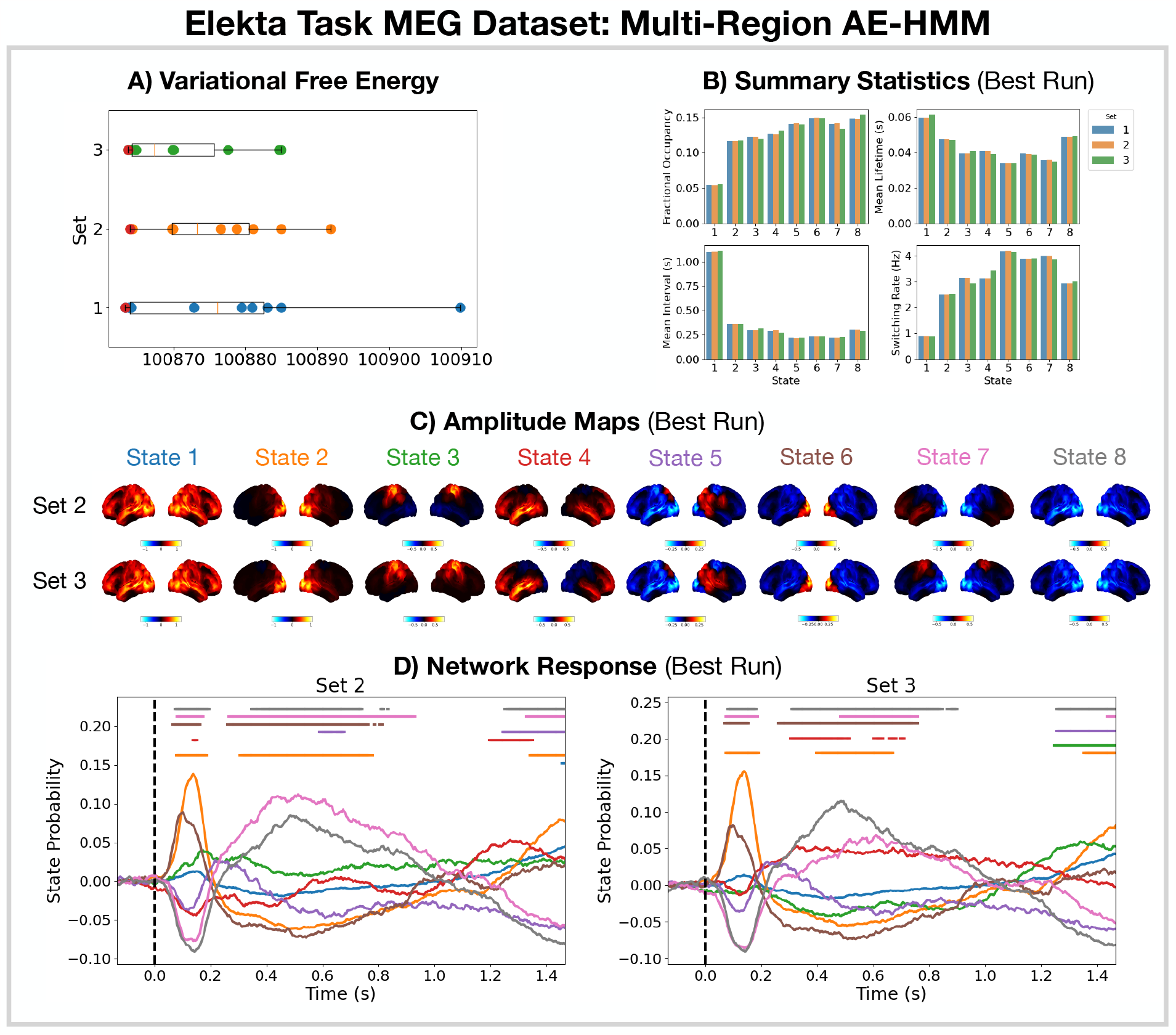
Reproducibility of the Elekta task MEG dataset multi-region AE-HMM analysis. A) Variational free energy for 3 sets of 10 runs. B) Summary statistics for the best run from each set (indicated by the red dot in A). C) Amplitude maps relative to the mean across states for the best run from sets 2 and 3. D) Network response to the visual task for sets 2 and 3. The horizontal bars indicate time points with p-values < 0.05. The maximum statistic pooling over states and time was used in permutation testing to control for the family-wise error rate.

**Figure S3:**
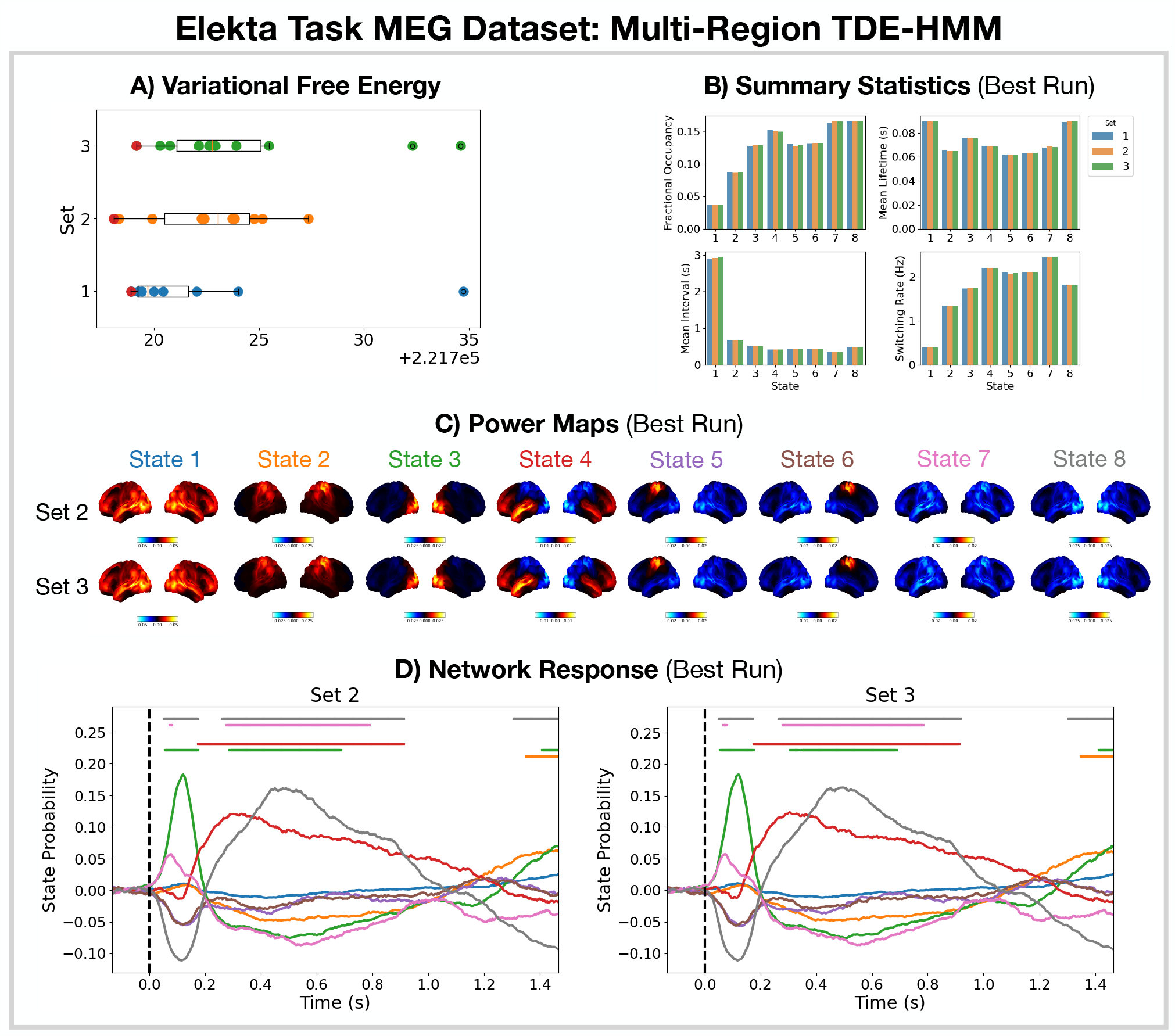
Reproducibility of the Elekta task MEG dataset multi-region TDE-HMM analysis. A) Variational free energy for 3 sets of 10 runs. B) Summary statistics for the best run from each set (indicated by the red dot in A). C) Power maps relative to the mean across states for the best run from sets 2 and 3. D) Network response to the visual task for sets 2 and 3. The horizontal bars indicate time points with p-values < 0.05. The maximum statistic pooling over states and time was used in permutation testing to control for the family-wise error rate.

**Figure S4:**
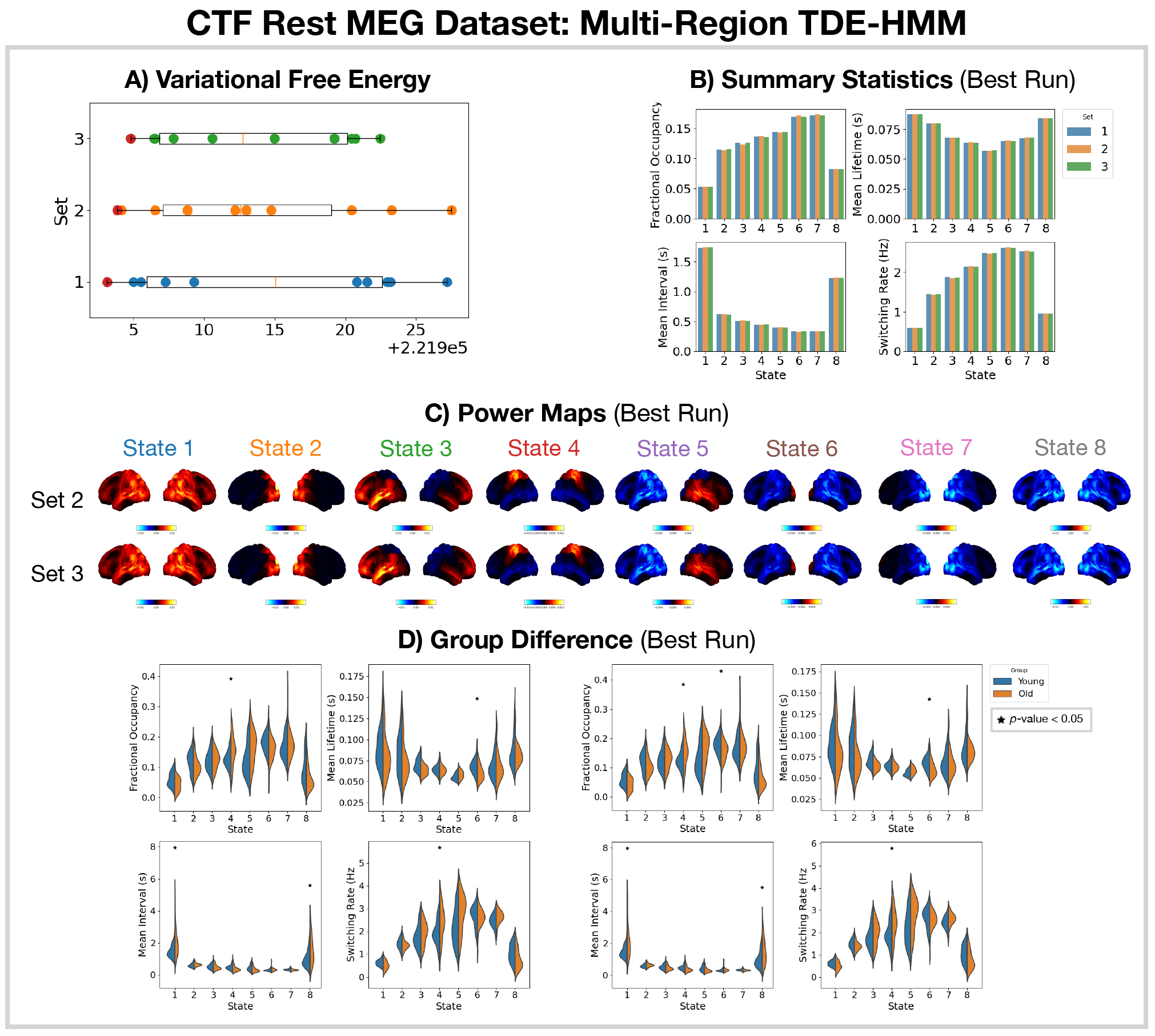
Reproducibility of the CTF rest MEG dataset multi-region TDE-HMM analysis. A) Variational free energy for 3 sets of 10 runs. B) Summary statistics for the best run from each set (indicated by the red dot in A). C) Power maps relative to the mean across states for the best run from sets 2 and 3. D) Comparison of the summary statistics for a young (18-34 years old) and old (34-60 years old) group. The star indicates a p-values < 0.05. The maximum statistic pooling over states and metrics was used to control for the family-wise error rate.

**Figure S5:**
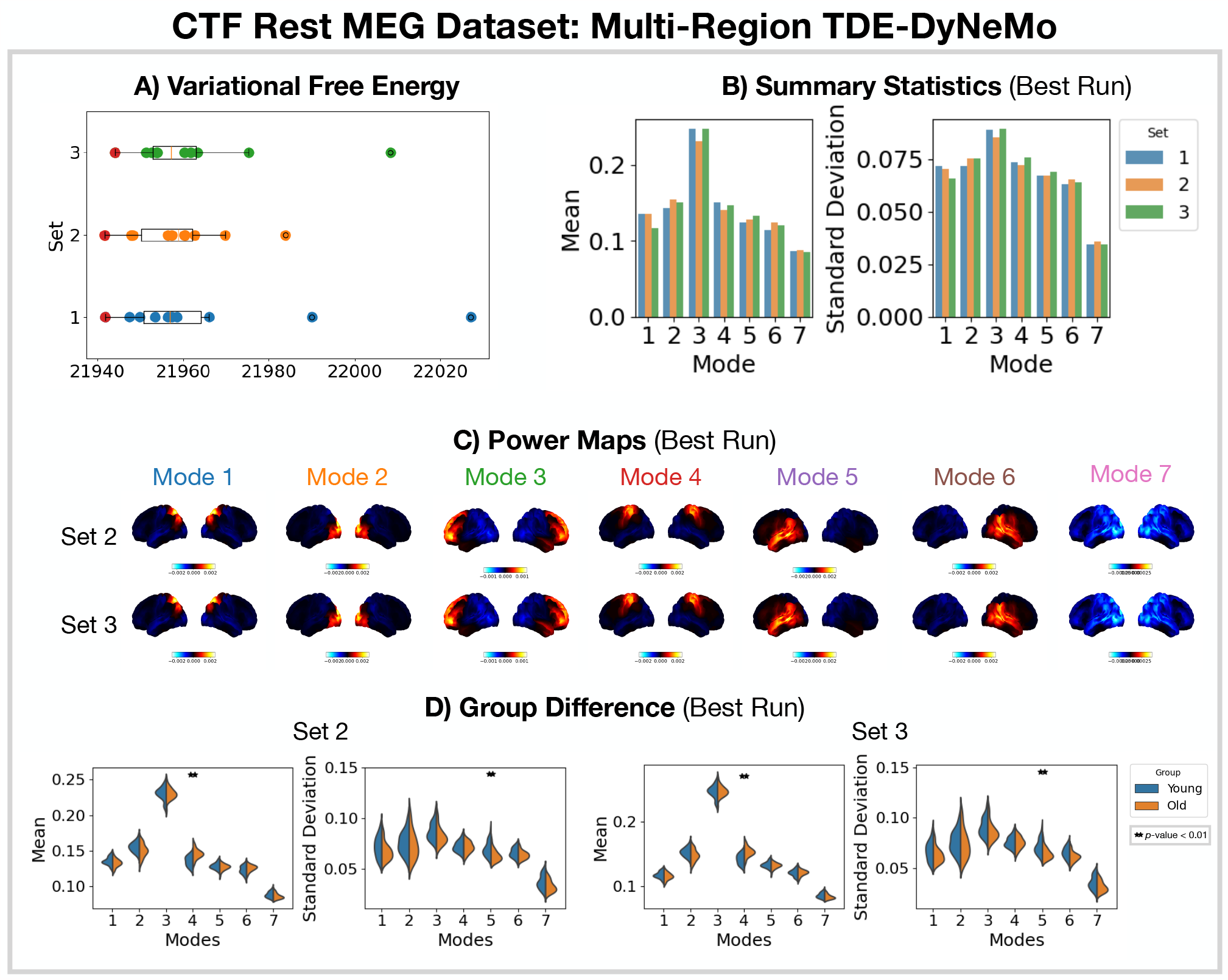
Reproducibility of the CTF rest MEG dataset multi-region TDE-DyNeMo analysis. A) Variational free energy for 3 sets of 10 runs. B) Summary statistics for the best run from each set (indicated by the red dot in A). C) Power maps relative to the mean across modes for the best run from sets 2 and 3. D) Comparison of the summary statistics for a young (18-34 years old) and old (34-60 years old) group. The double star indicates a p-values < 0.01. The maximum statistic pooling over modes and metrics was used to control for the family-wise error rate.

## Footnotes

1 This is equivalent to maximising the negative variational free energy, which is also known as the evidence lower bound (ELBO) [35].

2 Also known as the Viterbi path.

3 See `osl.preprocessing.osl wrappers.detect badsegments` and `detect badchannels` in [40].

4 See `osl.preprocessing.mne wrappers.run mne ica autoreject` in [40].

5 We used a parcellation based on anatomy: `fmri d100 parcellation with PCC reduced 2mm ss5mm ds8mm.nii.gz` [40].

6 See `osl.source_recon.sign_flipping` in [40]. Note, this step can be skipped if you are training on amplitude envelope data.

7 This step is analogous to dual regression in independent component analysis.

8 The lifetime is also known as *dwell time*.

9 We weight each mode time course by the trace of its mode covariance and divide by the sum over modes at each time point to ensure the renormalised mode time course sums to one.

10 A picture of a familiar, unfamiliar or scrambled face.

1 The full list of dependencies is given in the source code: osl-dynamics/setup.cfg.

